# A modular mechanism determines the proton-pumping stoichiometry of respiratory complex I

**DOI:** 10.64898/2026.10.08.757549

**Authors:** Robert A. Waddell, Harry L. Messent, Marcelo Mesa Costa-Lima, John J. Wright, Jack H. Salisbury, Judy Hirst

## Abstract

Respiratory complex I, a crucial metabolic enzyme, uses the reducing potential of NADH to pump protons across energy-transducing membranes, but its mechanism of catalysis remains unknown. Here, guided by detailed structural information, we combine targeted mutagenesis and biophysical analyses to define the proton-pumping pathways in the three antiporter-like (proton-pumping) subunits of complex I from *Paracoccus denitrificans* and *Escherichia coli*. In *Pd*-CI, a four-proton pump, one of the three proton-uptake pathways is inactive, reconciling the four-proton/three-subunit mismatch. In *Ec*-CI two pathways are inactive – and *Ec*-CI pumps only two protons. We define the structural determinants for redox-coupled proton uptake, establish a mechanistic framework for proton pumping, and reveal how the proton-pumping stoichiometry has been adapted to deliver efficient energy conservation in diverse bioenergetic environments.

## Introduction

Complex I (proton-pumping NADH:quinone oxidoreductase, CI), an energy-transducing molecular machine, is essential for oxidative phosphorylation in mammalian mitochondria and respiring organisms across the tree of life. It uses free energy from the reduction of quinone by NADH to pump protons across the mitochondrial inner or bacterial/archaeal cytoplasmic membrane and generate the proton-motive force (Δp) that drives ATP synthesis and transport processes. Although mammalian CI contains 45 protein subunits, only fourteen core subunits, which form the minimal prokaryotic enzyme and are conserved in an L-shaped structure, are necessary for catalysis (*1–4*). NADH is oxidized by a flavin mononucleotide at the top of the hydrophilic domain, then the electrons pass down a chain of iron-sulfur clusters to quinone, bound with its redox-active headgroup at the end of a long channel from the membrane. How the energy from redox catalysis is captured in the hydrophilic domain and tightly ‘coupled’ to proton pumping in the membrane domain (neither can occur independently) is disputed (*4–11*). Structural analyses have resolved membrane-domain hydrogen-bonding networks and structured waters in detail (*6*, *10*, *12–17*) but the ‘proton-wiring diagram’ that describes the proton uptake and output pathways, transfer connections and gates (control points) remains undefined. Without this underpinning knowledge of proton movement, the energy-transducing coupling mechanisms cannot be tackled effectively. Supplementary Figure 1 summarizes the fourteen-subunit core structure, and the key functional elements and residues investigated.

Mammalian CI pumps four protons from the negatively charged (N-side) to the positively charged (P-side) side of the membrane for every NADH oxidized and ubiquinone reduced (*18*). However, structures show the distal membrane domain of CI contains three homologous antiporter-like subunits (ND5, ND4, ND2), with candidate proton-uptake (N-side) and -output (P-side) pathways connected by a ‘central axis’ of essential charged residues (*6*, *10*, *12–17*, *19*) that couples redox catalysis to proton pumping (Supplementary Figure 1). The mismatch between the *four*-proton stoichiometry and *three* antiporter-like subunits has been challenging to reconcile in structure-based proposals for the catalytic mechanism, which are not distinguished by currently available functional and biophysical data. Proposals have included: four separate proton-transporter units, including a fourth uncharacterized unit in the proximal membrane domain (*7*, *8*, *11*); compensatory P-side proton uptake for quinone reduction (*9*); intersubunit proton exchange in an integrated multisubunit-transporter unit with shared uptake/output pathways (*4–6*, *10*, *20*); one or more subunits not contributing to proton uptake/output (*4–6*, *10*); *and* proton exchange between (in either direction) the antiporter-like subunits and redox catalysis (*4*– *6*, *10*).

Here, we reveal how a complex with *three* antiporter-like subunits pumps *four* protons during each catalytic cycle, and that the modular design has enabled different species to evolve different proton-pumping stoichiometries. Specifically, underpinned by detailed structural information (*6*, *14*), we deploy targeted mutagenesis in CI from *Paracoccus denitrificans* (*Pd*-CI), an α-proteobacterium with an aerobic, ubiquinone-10-dependent, mitochondrial-like respiratory chain and confirmed four-proton CI stoichiometry (*18*, *21–23*), and *Escherichia coli* (*Ec*-CI), a facultative anaerobic γ-proteobacterium that respires using both ubiquinone (UQ) and menaquinone (MQ) (*24*, *25*). We demonstrate that the proton-pumping stoichiometry is controlled by the number of redox-coupled proton-uptake pathways to enable efficient energy conservation under diverse bioenergetic conditions.

### Only subunits ND5 and ND4 contain redox-coupled proton-uptake pathways in *Pd*-CI

Figure 1A shows the proton-uptake pathways identified in subunits ND5, ND4 and ND2 of *Pd*-CI (*14*) using sequence conservation and structural information, including observed and simulated hydration patterns (*5*, *10*, *13*, *26*). Each pathway begins with a conserved ‘uptake-Lys’ at the N-side solvent-accessible surface, then descends to the central axis via a conserved uptake-Ser/Thr. The central axis enters each subunit via a water-bridged Glu-Lys ion pair, before meeting the uptake pathway (*6*, *10*, *12–14*, *17*) (Supplementary Figure 1). In ND5, they meet at a ‘junction-His’ on transmembrane helix (TMH) 8 that has been observed in two helical conformations, with different His-gated connectivities between the uptake pathway and upstream and downstream central axis (*16*, *27*, *28*). The proposed conformational interconversion is called the ND5 ‘master switch’. In ND4, the junction-His on TMH7, which has been observed in two rotameric conformations that similarly switch its connectivities (*16*), sits above a ‘central’ TMH8-Lys. ND2 is a close structural homologue of ND4, but its junction residue is Tyr in both *Pd*-CI and *Ec*-CI (*6*, *14*). After meeting the junction, the central axis crosses a subunit-dependent Lys/His/Tyr bridging path to a ‘terminal’ Lys/Glu, from which it either crosses to the next subunit (ND2/ND4) or meets the structurally defined P-side proton-output pathway (ND5) (*5*, *6*, *10*, *13*, *14*). Supplementary Figure 2 and Supplementary Table 1 summarize the conservation of key residues in all structurally characterized species of CI.

**Figure 1.**
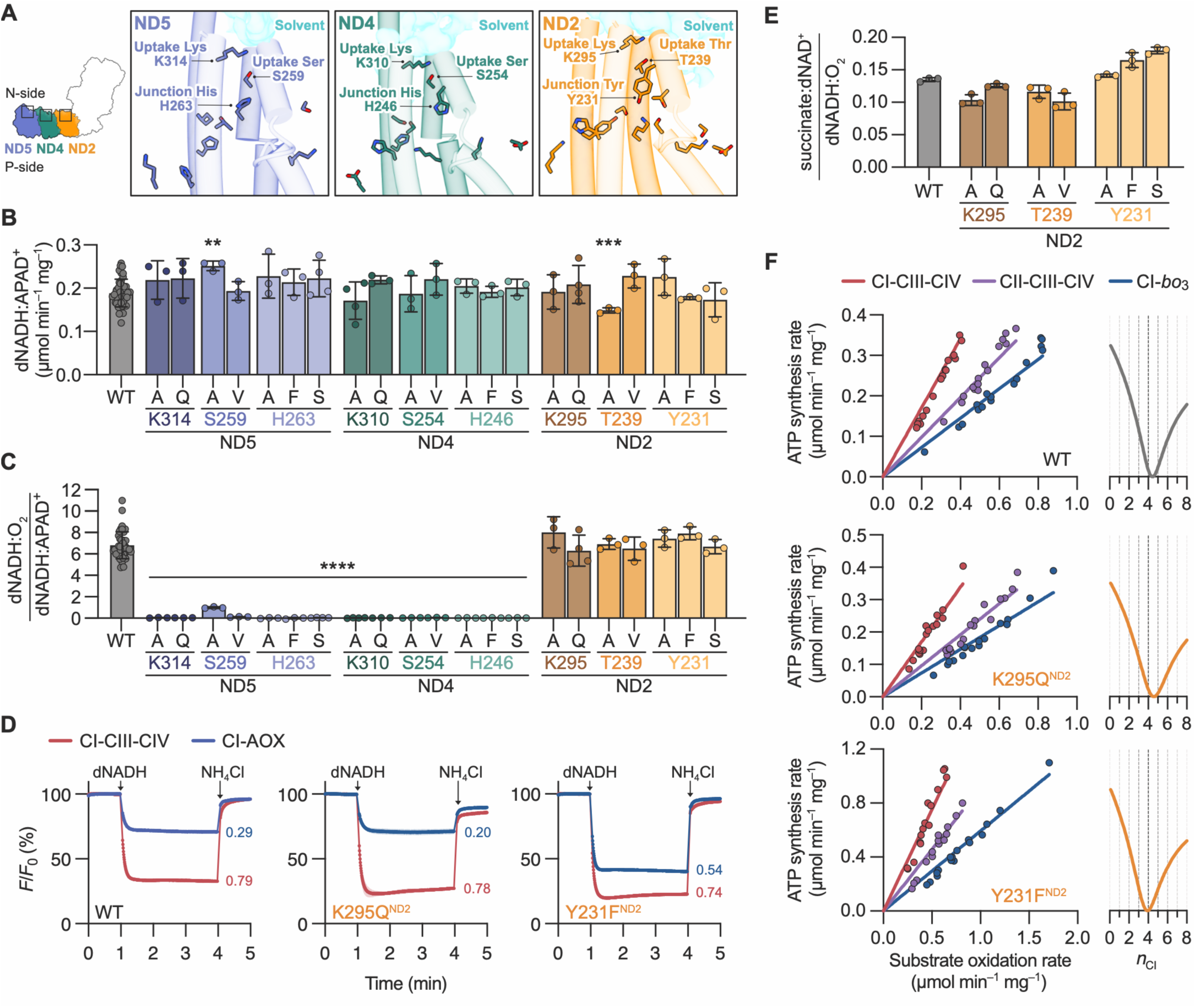
Redox catalysis is coupled to proton uptake in ND5 and ND4 but not ND2 in *P. denitrificans* complex. **I.** (**A**) The candidate proton-uptake pathways in *Pd*-ND5 (blue), *Pd*-ND4 (green) and *Pd*-ND2 (orange) showing the uptake-Lys, uptake-Ser/Thr, junction-His/Tyr and N-side solvent-accessible surfaces (PDB: 8QBY) (*14*). TMHs 7-8, 10-11 that house the pathways, and central axis residues are also shown (see also (*14*) and Supplementary Figure 1). (**B**) Rates of dNADH:APAD^+^ oxidoreduction by independent membrane preparations of *Pd*-CI variants (*N* ≥ 3, ± S.D.). (**C**) Normalized rates of *Pd*-CI turnover (piericidin A-sensitive dNADH:O_2_ (Supplementary Figure 3A-B) divided by dNADH:APAD^+^ oxidoreduction (panel B)) (*N* ≥ 3, ± S.D.). (**D**) Proton pumping in *Pd*-SBPs assessed by ACMA fluorescence quenching (mean average ± S.D. of three technical replicates). dNADH:O_2_ oxidoreduction by CI-CIII-CIV or CI-AOX was initiated by dNADH, then ΔpH dissipated by NH_4_Cl (see Supplementary Figure 5). Quenching is determined by both the proton-pumping stoichiometry (ten for CI-CIII-CIV, four for CI-AOX [for WT *Pd*-CI]) and the rate of dNADH oxidation (marked in μmol min^-1^ mg^-1^). Representative variants for the uptake-Lys (*Pd*-K295Q^ND2^) and junction-Tyr (*Pd*-Y231F^ND2^) are shown (see Supplementary Figure 6 for additional variants and analyses). (**E**) Reverse electron transfer (RET, succinate:dNAD^+^ oxidoreduction) rates in *Pd*-SBPs (*n* = 3 technical replicates, ± S.D.) normalized to dNADH:O_2_ oxidoreduction at Δp = 0 (see Supplementary Figure 7 for supporting data and information). (**F**) Determination of the proton-pumping stoichiometries of the *Pd*-WT, *Pd*-K295Q^ND2^ and *Pd*-Y231F^ND2^ variants (see Supplementary Figure 8 for further information). Left: ATP synthesis rates plotted against substrate oxidation rates for three pathways (CI-CIII-CIV dNADH:O_2_, CII-CIII-CIV succinate:O_2_, and CI-*bo*_3_ dNADH:O_2_ by *bo*_3_-proteoliposomes). The lines shown are the global best fits, and their relative gradients reflect the pathway stoichiometries (10, 6, 4 respectively, for *Pd*-WT). Right: The *Pd*-CI stoichiometry (*n*_CI_) is determined by plotting the normalized log of the sum of the squared errors (data vs. fit) at each candidate value of *n*_CI_. The global best fit is at the minimum point (*Pd*-WT = 4.44, *R*^2^ = 0.923; *Pd*-K295Q^ND2^ = 4.59, *R*^2^ = 0.911; *Pd*-Y231F^ND2^ = 3.90, *R*^2^ = 0.941) and the y-scale is from 1 to 100. See Materials and Methods for experimental conditions and protocols.

To investigate the activity of each proposed proton-uptake pathway during catalysis we used unmarked mutagenesis in the *P. denitrificans* genome (*18*, *22*, *23*). Figure 1B shows the rates of dNADH:APAD^+^ oxidoreduction in membrane preparations from a set of uptake-pathway variants. Deamino-NADH (dNADH) is CI-specific (it is not oxidized by the NDH-2 NADH:quinone oxidoreductase introduced to compensate for inactive CI) and so the flavin-localized dNADH:APAD^+^ reaction (APAD^+^ is an NAD^+^ analogue) reports on the CI abundance (*22*) – which remains consistent in all variants. In contrast, rates of dNADH:O_2_ oxidoreduction (catalyzed by CI-CIII-CIV and sensitive to the CI-inhibitor piericidin A, Supplementary Figure 3A-B) vary substantially from wild-type (WT)-like activity to inactive; they were normalized by the dNADH:APAD^+^ rates to evaluate the catalytic competence of each variant, independent of its expression level (Figure 1C). Strikingly, mutations in the ND4 and ND5 proton-uptake pathways stop catalysis completely, but equivalent ND2 mutations have no effect.

Each junction residue (*Pd*-H263^ND5^, *Pd*-H246^ND4^, *Pd*-Y231^ND2^) was mutated to Ala, Phe and Ser. The inactive ND5 and ND4 variants (*vs*. the fully active ND2 variants) (Figure 1C) support the unique ability of a junction-His to gate proton transfer (by forming separate interactions at Nδ and Nε) being crucial in these subunits (*16*, *27*, *28*). To confirm that mutagenesis in ND2 is able to stop catalysis, all three ion-pair Lys-to-Gln variants were shown to be inactive (Supplementary Figure 3). Each uptake-Ser/Thr (*Pd*-S259^ND5^, *Pd*-S254^ND4^, *Pd*-T239^ND2^) was mutated to Ala and Val, and each uptake-Lys (*Pd*-K314^ND5^, *Pd*-K310^ND4^*, Pd*-K295^ND2^) to Ala and Gln. The inactive ND5 and ND4 variants (*vs*. the fully-active ND2 variants) (Figure 1C) further supports proton uptake only in ND5 and ND4 during catalysis. Although the minimal activity of *Pd*-S259A^ND5^ suggests the uptake-Ser γ-hydroxyl can be poorly substituted by an intercalated water, the inactive *Pd*-K314^ND5^ and *Pd*-K310^ND4^ variants show the uptake-Lys sidechains cannot be substituted by unstructured waters, consistent with suggestions that, in ND5, the uptake-Lys influences the junction-His conformation (*27*, *28*). Finally, the isolated *Pd*-K314Q^ND5^ and *Pd*-K310Q^ND4^ complexes were homogeneous, intact and stable (Supplementary Figure 4). Therefore, the ND5/ND4 uptake-Ser and uptake-Lys mutations, which stop catalysis from outside the central axis, demonstrate that *Pd*-K314-S259-H263^ND5^ and *Pd*-K310-S254-H246^ND4^ form active redox-coupled proton-uptake pathways. Contrary to previous proposals (*6*), they cannot compensate for one another so there is no common pool of protons between them.

Although our results strongly indicate *Pd*-ND2 does not uptake protons, we further considered the alternative possibility that the mutations could be both blocking and decoupling an active pathway, thus allowing redox catalysis to continue unhindered. As decoupling the pathway would decrease the proton-pumping stoichiometry, we used inverted membrane vesicles (sub-bacterial particles, SBPs (*18*, *23*)) to test proton pumping in the *Pd*-ND2 variants. First, in ACMA proton-pumping assays (*22*, *29–31*), which report on the formation of ΔpH across the SBP membrane (Supplementary Figure 5), all the variants displayed WT-like behavior (Figure 1D and Supplementary Figure 6). Second, all the variants were able to perform reverse electron transfer (RET, Δp-driven dNAD^+^:ubiquinol oxidoreduction (*23*, *30*)), which relies on the combined free energy of all four pumped protons (Figure 1E and Supplementary Figure 7). Finally, like WT *Pd*-CI (*18*, *23*), the *Pd*-K295Q^ND2^ and *Pd*-Y231F^ND2^ variants each display a four-proton stoichiometry (Figure 1F and Supplementary Figure 8). Therefore, *Pd*-K295-T239-Y231^ND2^ does not form an active redox-coupled proton-uptake pathway: subunit ND2 does not take up protons during catalysis.

### Junction-His residues couple proton uptake to redox catalysis

Junction-Tyr^ND2^ lacks the Nδ and Nε centers of the junction-His^ND5/ND4^, which form separate interactions to gate proton-transfer reactions (Figure 2A) (*16*, *27*, *28*), suggesting the junction-residue identity determines the uptake activity. Therefore, we exchanged the junction-His and junction-Tyr. *Pd*-H263Y^ND5^ is inactive, confirming the junction-His^ND5^ is essential (Figure 2B). *Pd*-Y231H^ND2^ showed WT-like behavior in dNADH oxidation, RET and proton pumping ACMA assays (Figure 2B-C, Supplementary Figure 9D), and its proton-stoichiometry is four (Figure 2E). Therefore, introducing a junction-His^ND2^ does not activate redox-coupled proton uptake in ND2. Consistent with the opposing behaviors of the Ala/Phe/Ser variants (inactive in ND4, active in ND2, Figure 1C), additional factors must also determine pathway function, and structure comparisons highlight the route of the central axis (*6*, *10–14*, *17*) (Supplementary Figure 1). In *Pd*-ND5 the central axis rises to ‘peak’ at the junction-His (Supplementary Figure 9E), and in *Pd*-ND4 we propose it does likewise (Figure 2F), triggering uptake in each case (*16*). Conversely, in *Pd*-ND2 the junction-Tyr hydroxyl reaches down, suggesting a ‘flat’ route that includes the central-Lys (*Pd*-K247^ND2^), which sits below the junction in both ND4 and ND2 (Figure 2F). *Pd*-K247Q^ND2^ catalyzes with decreased dNADH-oxidation rate, but WT-like characteristics in RET and ACMA assays (Figure 2B-C, Supplementary Figure 9D), consistent with redundancy in the central-axis network (*14*). In contrast, hydrophobic residues obstruct the flat route in *Pd*-ND4 and the central-Lys^ND4^ faces away from the junction-His^ND4^, both promoting the peak route (Figure 2F). Finally, *Pd*-H246Y^ND4^ is partially active in dNADH-oxidation assays but unable to catalyze RET, displays decreased proton pumping relative to redox catalysis in CI-linked ACMA assays, and has a decreased proton-pumping stoichiometry (Figure 2B-E), indicating the junction is dysregulated. The nearby *Pd*-K263Q^ND4^ central-Lys variant displays very low activity in dNADH-oxidation assays (Figure 2B) and CI-linked ACMA assays (Supplementary Figure 9D) suggesting it is also dysregulated, but the low rates preclude conclusive analyses (Supplementary Figure 7). We thus propose three requirements for redox-coupled proton uptake: a competent proton-uptake pathway; a junction-His that gates the uptake pathway/central axis intersection; *and* structures that confine the central axis to engage the junction-His to trigger uptake.

**Figure 2.**
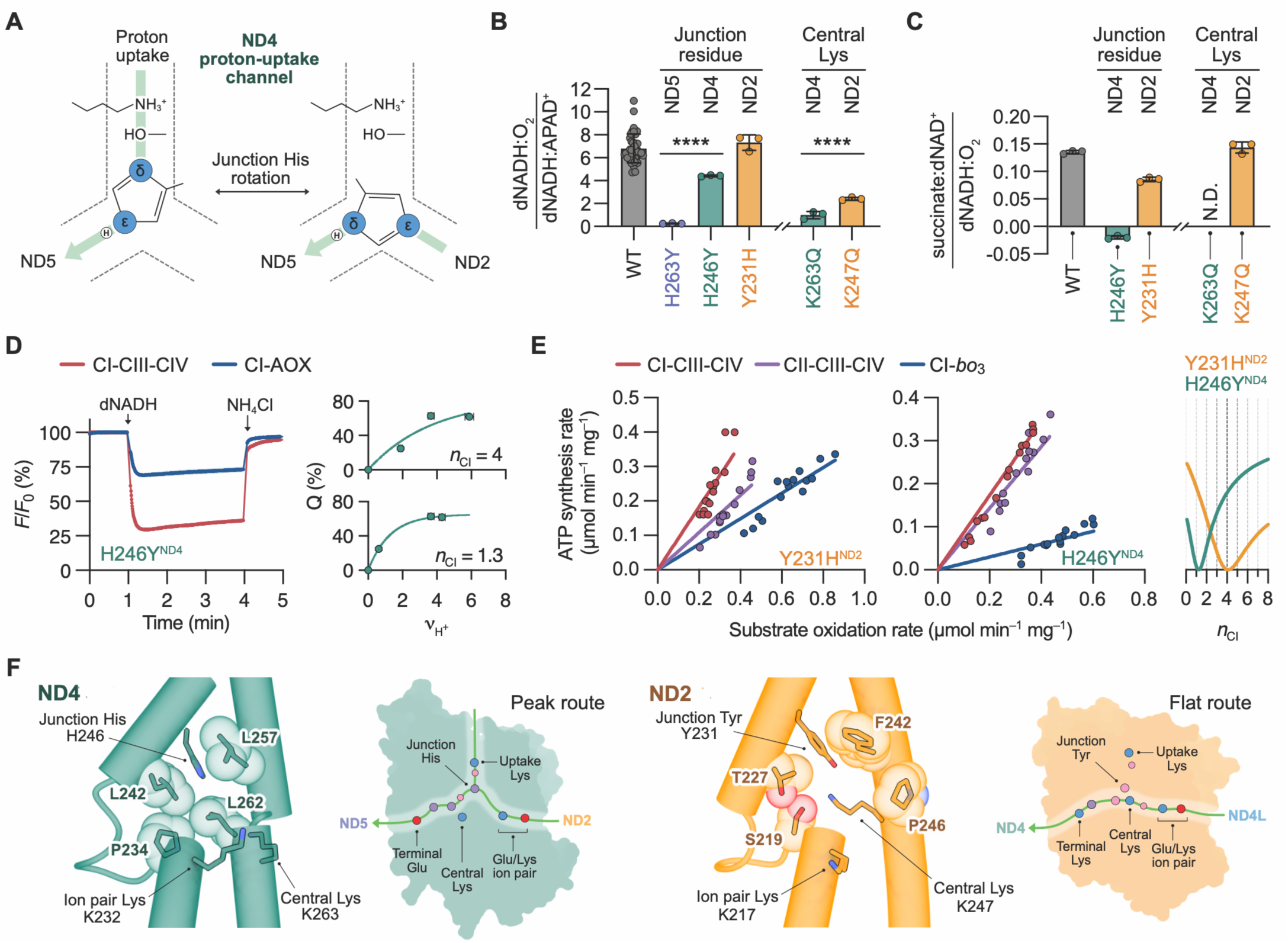
Histidine junctions couple proton uptake to redox catalysis in *Pd*-CI. (**A**) Schematic diagram showing the junction-His^ND4^ switching conformation to enable Nδ and Nε to catalyze proton transfer along different pathways (left, uptake pathway to downstream central axis; right, upstream to downstream central axis) (*16*). (**B**) Normalized rates of *Pd*-CI turnover (piericidin A-sensitive dNADH:O_2_ divided by dNADH:APAD^+^ oxidoreduction, Supplementary Figure 9A-C), *N* ≥ 3, ± S.D.). (**C**) Reverse electron transfer (RET, succinate:dNAD^+^ oxidoreduction) rates in *Pd*-SBPs (*n* = 3 technical replicates, ± S.D.) normalized to the rate of dNADH:O_2_ oxidoreduction at Δp = 0 (see Supplementary Figure 7 for the separate rates and further information). (**D**) Proton pumping in *Pd*-H246Y^ND4^ SBPs assessed by ACMA fluorescence quenching (mean average ± S.D. of three technical replicates). dNADH:O_2_ oxidoreduction by CI-CIII-CIV or CI-AOX was initated by dNADH, then ΔpH dissipated by NH_4_Cl. ACMA analyses of four different pathways (see Supplementary Figure 5) are not consistent with *n*_CI_ = 4 but are consistent with *n*_CI_ = 1.3 (the best-fit value from panel E). See Supplementary Figure 9D for additional variants and analyses. (**E**) Determination of the proton-pumping stoichiometries of the *Pd*-Y231H^ND2^ and *Pd*-H246Y^ND4^ variants (see Supplementary Figure 8 for further information). Left: ATP synthesis rates plotted against substrate oxidation rates for three pathways (dNADH:O_2_ by CI-CIII-CIV, succinate:O_2_ by CII-CIII-CIV, and dNADH:O_2_ by CI-*bo*_3_ using *bo*_3_-proteoliposomes). The lines shown are the global best fits, and their relative gradients reflect the pathway stoichiometries. Right: The *Pd*-CI stoichiometry (*n*_CI_) is determined by plotting the normalized log of the sum of the squared errors (data vs. fit) at each candidate value of *n*_CI_. The global best fit is at the minimum point (*Pd*-Y231H^ND2^ = 4.11, *R*^2^ = 0.793; *Pd*-H246Y^ND4^ = 1.25, *R*^2^ = 0.942) and the y-scale is from 1 to 100. *Pd*-Y231H^ND2^ matches WT-CI but the stoichiometry of *Pd*-H246Y^ND4^ is decreased. See Materials and Methods for experimental conditions and protocols. (**F**) Structural and cartoon representations of the peak and flat routes of the central axis in *Pd*-ND4 and *Pd*-ND2, respectively. The view along the central axis from the ion pair (structures) shows TMH7-8 with van der Waals radii for relevant residues. The flat pathway is blocked in *Pd*-ND4 but clear in *Pd*-ND2. The cartoons show the profile of the central axis and uptake pathway viewed from the membrane (red – Glu, blue – Lys, purple – His, pink –Ser/Tyr/Thr). See Supplementary Figure 9E for equivalent information on ND5.

Molecular dynamics simulations have proposed that three conserved N-side Phe, adjacent to the junction residues (Figure 3A, Supplementary Figure 1), gate proton uptake by moving into and out of the pathways, alternately breaking and reforming water networks and proton-transfer connectivity (*7*, *10*, *26*, *32*). Therefore, each Phe (*Pd*-F355^ND5^, *Pd*-F354^ND4^, *Pd*-F339^ND2^) was changed to Ala and Leu. In dNADH-oxidation assays, both ND5 variants and *Pd*-F354A^ND4^ were partially active, whereas both ND2 variants and *Pd*-F354L^ND4^ displayed WT-like activity (Figure 3B). *Pd*-F355A^ND5^ and *Pd*-F339A^ND2^ displayed WT-like behavior in RET and proton-pumping ACMA assays, and four-proton stoichiometries (Figure 3C-D, Supplementary Figures 10-11), excluding the Phe as a proton-uptake gate in ND5 (and ND2). The same was true for F354L^ND4^, but *Pd*-F354A^ND4^, like adjacent H246Y^ND4^, showed dysregulated behavior (decreased relative ACMA fluorescence quenching, unable to catalyze RET, decreased stoichiometry) (Figure 3C-D, Supplementary Figure 10-11). Structural analyses (*16*) suggested the bulky Phe corrals the junction-His^ND4^ as it switches conformation (Figure 3A, E), consistent with a common mode of dysfunction affecting both H246Y^ND4^ and F354A^ND4^. However, the double *Pd*-F354A:K310Q^ND4^ variant is inactive (Figure 3B) so dysregulated catalysis still relies on the uptake-Lys (*16*, *27*, *28*): unlike in ND2, the pathway in *Pd*-F354A^ND4^ is not decoupled from redox catalysis, and the dysregulation likely reflects failure to execute a specific coupling step. Dysregulation of the delicate rotameric ND4 junction contrasts with the robust conformational master-switch ND5 junction, for which decreased stoichiometry variants have not been identified. Finally, the Phe-gating mechanism was also proposed for two P-side residues (*Pd*-F406^ND4^, *Pd*-F392^ND2^) at the start of two proposed proton-output pathways (*8*, *26*, *32*). However, both Phe-to-Ala variants exhibited WT-like behavior in dNADH-oxidation, ACMA and RET assays, along with four-proton stoichiometries (Figure 3B-C, Supplementary Figures 10-11). We conclude that proton uptake in ND5 and ND4 is gated by junction-His conformation, not by hydrophobic residues that control pathway hydration.

**Figure 3.**
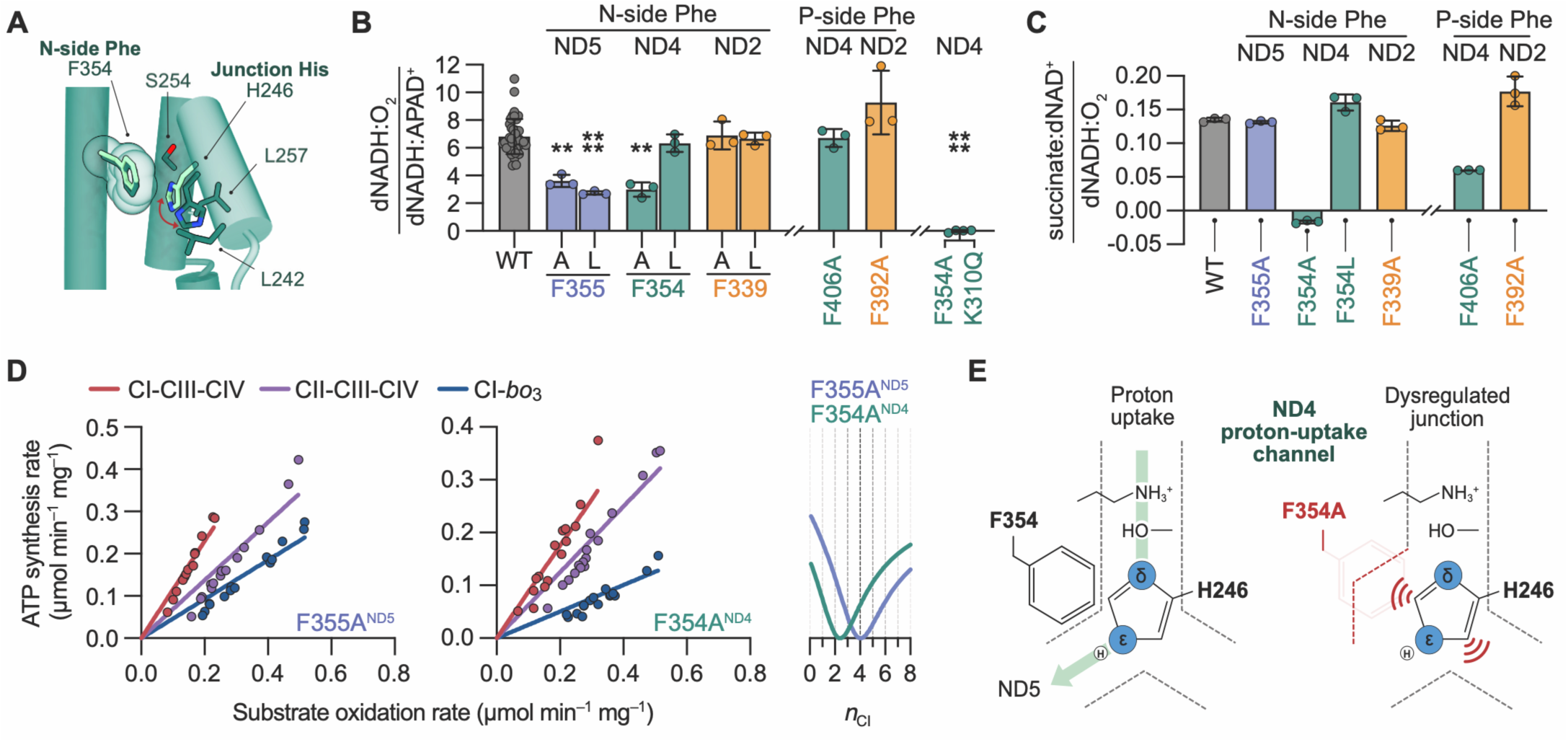
Phenylalanine residues shape channel architecture but do not gate proton transfer. (**A**) The N-side phenylalanine constrains the ND4 junction-His to constructive conformations for proton transfer. The structure of *Pd-*CI (PDB: 8QBY) (*14*) is shown in dark green with *Pd*-H246^ND4^ in the ‘planar’ state between *Pd*-L242^ND4^ and *Pd*-L257^ND4^. The junction-His and N-side Phe from the ‘perpendicular’ state of *B. taurus* CI (PDB: 9R12) (*16*) (light green) are superimposed. (**B**) Normalized rates of *Pd*-CI turnover (piericidin A-sensitive dNADH:O_2_ divided by dNADH:APAD^+^ oxidoreduction, Supplementary Figure 10A-C), *N* ≥ 3, ± S.D.) for N- and P-side Phe variants. F354:K310^ND4^ is a double variant. (**C**) Reverse electron transfer (RET, succinate:dNAD^+^ oxidoreduction) rates in *Pd*-SBPs (*n* = 3 technical replicates, ± S.D.) for N- and P-side Phe variants normalized to the rate of dNADH:O_2_ oxidoreduction at Δp = 0 (see Supplementary Figure 7 for the separate rates and further information). (**D**) Determination of the proton-pumping stoichiometries of the *Pd*-F355A^ND5^ and *Pd*-F354A^ND4^ variants of *Pd*-CI (see Supplementary Figure 8 for further information). Data on the additional N- and P-side Phe variants are in Supplementary Figure 10D. Left: ATP synthesis rates plotted against substrate oxidation rates for three pathways (dNADH:O_2_ by CI-CIII-CIV, succinate:O_2_ by CII-CIII-CIV, and dNADH:O_2_ by CI-*bo*_3_ using *bo*_3_-proteoliposomes). The lines shown are the global best fits, and their relative gradients reflect the pathway stoichiometries. Right: The *Pd*-CI stoichiometry (*n*_CI_) is determined by plotting the normalized log of the sum of the squared errors (data vs. fit) at each candidate value of *n*_CI_. The global best fit is at the minimum point (*Pd*-F355A^ND5^ = 4.04, *R*^2^ = 0.894; *Pd*-F354A^ND4^ = 2.42, *R*^2^ = 0.906) and the y-scale is from 1 to 100. *Pd*-F255A^ND5^ matches WT-CI but the stoichiometry of *Pd*-F354A^ND4^ is decreased. See Materials and Methods for experimental conditions and protocols. (**E**) Schematic diagram of the ND4 proton-uptake pathway in *Pd*-WT and *Pd*-F354A^ND4^. Removing the Phe sidechain introduces conformational freedom and decreases stabilization of the productive proton-uptake conformation of the junction-His.

### Subunit ND4 in menaquinone-reducing complex I does not catalyze proton uptake

The uptake-pathway triads in mammalian and *P. denitrificans* ND5 and ND4 are identical, but in mammalian ND2 Arg-Gly-Thr replaces *Pd*-Lys-Thr-Tyr. Thus, consistent with their shared four-proton stoichiometry (*18*), redox-coupled proton uptake in mammalian-CI occurs only in ND5 and ND4. Wider analyses of all the structurally defined complexes (from animals, plants, protists, single-cell eukaryotes and bacteria) (Supplementary Table 1 and Supplementary Figure 2) further revealed universal conservation of the essential ND5-uptake triad, but diverse substitutions in the inactive ND2-uptake triad. Strikingly, the ND4-uptake triad is widely conserved, but missing in *Mycobacterium smegmatis* (Met-Ala-Ala replaces *Pd*-Lys-Ser-His) (*33*), indicating *Ms-*ND4 does not uptake protons. *Ms*-CI reduces MQ, and NADH:MQ oxidoreduction yields less free energy than NADH:UQ (ΔE_m,7_ ∼0.26 V *vs*. ∼0.39 V (*34*)), so we hypothesize that *Ms-*CI balances the energy equation by pumping fewer protons, enabling MQ-catalysis against high Δp (*35*).

Two further structurally characterized species also use MQ: *Thermus thermophilus* (MQ-dependent (*36*)) and *E. coli* (UQ or MQ, depending on the conditions (*24*, *25*)). In both, an uptake-Asp^ND4^, which replaces the uptake-Ser^ND4^ (*Tt*-H218-D228-K282^ND4^, *Ec*-H248-D258-K312^ND4^, Figure 4A and Supplementary Table 1) (*6*, *37*), is positioned in the pathway by the altered structure of ND4-TMH8 and interacts with the junction-His, perhaps restricting its movement (*16*). ND4-TMH8 also rotates the *Ec*-ND4 central-Lys from its *Pd*-ND4 position towards its *Pd*-ND2 position and clears bulky residues from the central-axis flat route (Figures 4A, 2F). These observations question whether *Ec*-ND4 and *Tt*-ND4 uptake protons and suggest the proton-pumping stoichiometries (*18*) may be altered.

**Figure 4.**
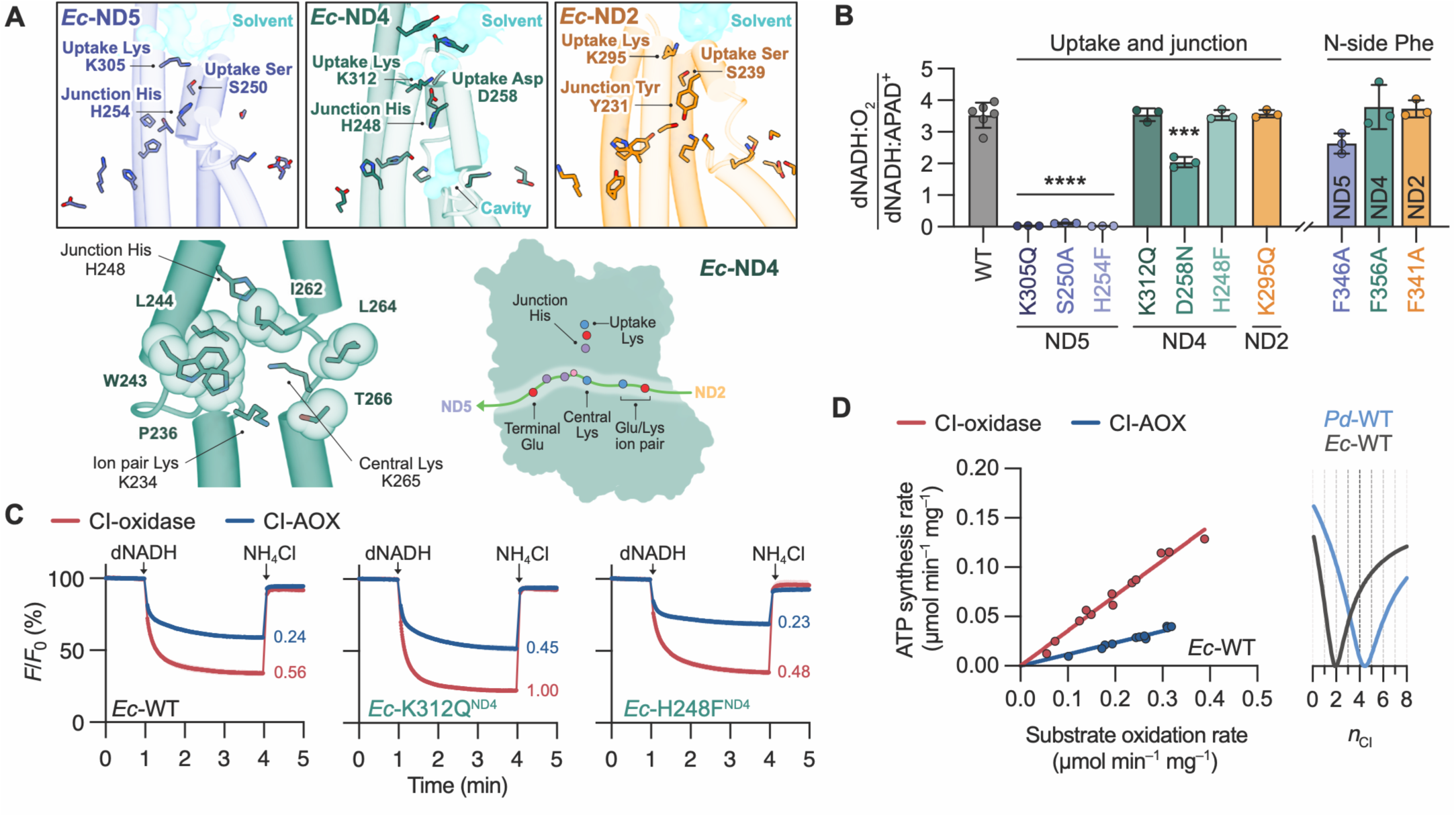
*E. coli* complex I couples redox-catalysis to proton uptake through ND5 (but not ND4 or ND2) with a proton-pumping stoichiometry of two protons per NADH oxidized. (**A**) Top: the candidate proton-uptake pathways in *Ec*-ND5 (blue), *Ec*-ND4 (green) and *Ec*-ND2 (orange) showing the uptake-Lys, uptake-Ser/Asp, junction-His/Tyr and N-side solvent-accessible surfaces (PDB: 7Z7S) (*6*). TMHs 7-8, 10-11 that house the pathways, and central axis residues are also shown. Bottom: structural and cartoon representations of the flat route of the central axis in *Ec*-ND4. The view along the central axis from the ion pair shows TMH7-8 with van der Waals radii for relevant residues. The cartoon shows the flat profile of the central axis and uptake pathway viewed from the membrane (red – Glu/Asp, blue – Lys, purple – His, pink –Ser). See Figure 2F for equivalent information on *Pd*-ND4. (**B**) Normalized rates of *Ec*-CI turnover (piericidin A-sensitive dNADH:O_2_ divided by dNADH:APAD^+^ oxidoreduction, Supplementary Figure 13A-C) determined in independent membrane preparations of *Ec*-CI variants, *N* ≥ 3, ± S.D. (**C**) Proton pumping in *Ec*-SBPs assessed by ACMA fluorescence quenching (mean average ± S.D. of three technical replicates). See Supplementary Figure 13D for supporting data on rates of *Ec*-SBP catalysis. dNADH:O_2_ oxidoreduction by CI-CIII-CIV or CI-AOX was initiated by dNADH, then ΔpH dissipated by NH_4_Cl. Quenching is determined by both the proton-pumping stoichiometry and the rate of dNADH oxidation (marked in μmol min^-1^ mg^-1^). Representative variants for the uptake-Lys (*Ec*-K312Q^ND4^) and junction-His (*Ec*-H248F^ND4^) are shown. (**D**) Determination of the proton-pumping stoichiometry of *Ec*-CI using the parental strain. Left: ATP synthesis rates plotted against substrate oxidation rates for two pathways (dNADH:O_2_ by CI-*bo*_3_ oxidase (native) and CI-AOX using AOX-proteoliposomes). The lines shown are the global best fit, and their relative gradients reflect the pathway stoichiometries. Right: The *Ec*-CI stoichiometry (*n*_CI_) is determined by plotting the normalized log of the sum of the squared errors (data vs. fit) at each candidate value of *n*_CI_. The global best fit is at the minimum point (1.96, *R*^2^ = 0.984) and the y-scale is from 1 to 100. The data from *Ec*-CI is shown relative to data from WT *Pd*-CI (Figure 1). Supporting information on the method and replicate measurements on different *Ec-*SBP preparations are given in Supplementary Figure 14. Only two respiratory pathways were accessible for ACMA experiments and stoichiometry measurements on *Ec*-SBPs precluding the more extensive three or four-pathway analysis applied to *Pd*-SPBs.

To investigate the uptake pathways in *Ec*-CI, we established a Δ*nuo E. coli* strain with the *nuo* operon expressed on an inducible plasmid (Supplementary Figure 12) (*38*). First, we mutated the three uptake-Lys (Figure 4B). *Ec*-K305Q^ND5^ is inactive whereas *Ec*-K295Q^ND2^ displays WT-like activity, confirming the ND5-uptake pathway is catalytically active and the ND2-uptake pathway is inactive. Strikingly, *Ec*-K312Q^ND4^ also displays WT-like activity, indicating that, in *Ec*-CI, the ND4-uptake pathway is also inactive. The junction-His *Ec*-H248F^ND4^ variant also displayed WT-like activity (the corresponding *Ec*-H254F^ND5^ was inactive), and both *Ec*-K312Q^ND4^ and *Ec*-H248F^ND4^ were confirmed to pump protons in ACMA assays using *E. coli* SBPs (Figure 4C). The uptake-Asp *Ec*-D258N^ND4^ variant displayed moderately decreased activity, perhaps because it introduces conformational freedom to the junction-His^ND4^ (Figure 4B). For the N-side Phe, *Ec*-F356A^ND4^ and *Ec*-F341A^ND2^ displayed WT-like activity, whereas *Ec*-F346A^ND5^ was partially active. We conclude the proton-uptake pathways in *Ec*-ND4 and *Ec*-ND2 are inactive: only *Ec*-ND5 catalyzes redox-coupled proton uptake. Finally, we determined the stoichiometry of *Ec*-CI by extending our stoichiometry protocol to *Ec*-SBPs from the parental strain, and using the four-proton stoichiometry of cytochrome *bo*_3_ oxidase (*39*, *40*) as a standard (Supplementary Figure 14). Our data (Figure 4D) indicate that *Ec*-CI pumps two protons (not four) for each NADH oxidized.

## Discussion

In canonical four-proton CI, including mammalian-and *Pd*-CI, the ND5 and ND4 proton-uptake pathways are redox-coupled and active, but the ND2 pathway is not ([++-]CI). In contrast, *E. coli* CI is a two-proton [+--]CI. Thus, there is no universal CI stoichiometry. Analogous to varying the *c*-ring in ATP-synthase (*41*, *42*), evolution has exploited the modular architecture of the proton-pumping domain and controlled the activities of the proton-uptake pathways to adapt the proton stoichiometry to different bioenergetic landscapes, demonstrated here for UQ-and MQ-based respiration. We suggest that CI-superfamily stoichiometries evolved initially by conserving the MrpA(ND5)-type subunit of the ancestral sodium-proton antiporter and increasing the number of MrpD(ND4/ND2)-type subunits (*43–45*), then, following mechanistic advances that increased the per-subunit proton yield, by silencing specific MrpD-type subunits by decoupling (and eliminating) their uptake pathways. Specifically, [++-]CI evolved to exploit the free-energy abundance of aerobic respiration with high-potential quinones at (or slightly before) the great oxidation event (*46*, *47*), then [+--]CI evolved later to utilize alternative terminal electron acceptors in anaerobic environments (*48–50*). Robust quantitative data on expanded sets of both native and engineered variants are now needed to further advance this integrated mechanistic, bioenergetic and evolutionary picture.

Four-proton [++-]CI excludes mechanisms in which four separate pathways each pump one proton (*7*, *8*, *11*), or three subunits each pump two, compensated by two P-side protons (*9*). Here, we discuss three models consistent with the balance of extant experimental data, which can each be adapted to both [++-] and [+--]CI (Figure 5). All three rely on intersubunit proton transfer along the central axis (*4*, *6*, *10*, *20*), a single proton-output pathway in ND5 (*5*, *6*, *10*, *13*, *14*), and the ability of the junction-His^ND5^ master switch to gate proton transfer from the upstream/ND4 central axis to the ND5-output pathway by a mechanism only partially substantiated by structural data (*16*). The models differ in mode of energy transfer along the central axis, from redox catalysis to the proton-pumping machinery: model-A, direct energy transfer, perhaps mediated by electrostatics (*7*, *51*); model-B, transfer of high-energy protons from redox catalysis (*4*, *10*); or model-C proton abstraction by redox catalysis creating high-energy proton holes (*5*, *6*). In model-A, ND4 and ND5 in [++-]CI each uptake two protons and all four exit from ND5, requiring alternating gating by the ND5-master switch (uptake-to-output, ND4-to-output) – whereas in [+--]CI it must catalyze differently, only uptake-to-output. In model-B, the junction-His^ND4^ catalyzes alternating gating in [++-]CI, using a ‘clockwork’ mechanism underpinned by two structurally defined states (*16*), but is bypassed in [+--]CI, as described above. The ND5-master switch similarly catalyzes alternating gating, but by a common mechanism, cycling twice in [++-]CI and once in [+--]CI. This single-stroke mechanism provides a clear proton-counting strategy, but how one extreme-energy proton is generated, transferred and utilized without energy loss is so-far unknown. In model-C, proton abstraction must occur by ND5-uptake (assuming a conserved mechanism) but in [++-]CI the abstracted (ND5-to-ND4) and pumped (ND4-to-ND5) protons must cross, and the ND5-master switch gate three different proton trajectories – although high-energy proton abstraction through ND5 offers an attractive explanation for redox-coupled CO_2_ formation in carbon-concentrating NDH-1 (*52*, *53*). In summary, by defining proton-uptake pathways and junction gates, we have imposed boundaries on the CI mechanism, distinguishing existing proposals, and highlighting the most fundamental unanswered questions: how energy is captured by redox catalysis and transferred to the proton-pumping domain. Uniting information from CI variants and the CI superfamily, where variations in subunit modularity, reaction thermodynamics and proton-pumping stoichiometry exist within a common architecture, now offers a powerful approach to transform understanding of these proton-pumping modular biomachines that deliver the energy of life.

**Figure 5.**
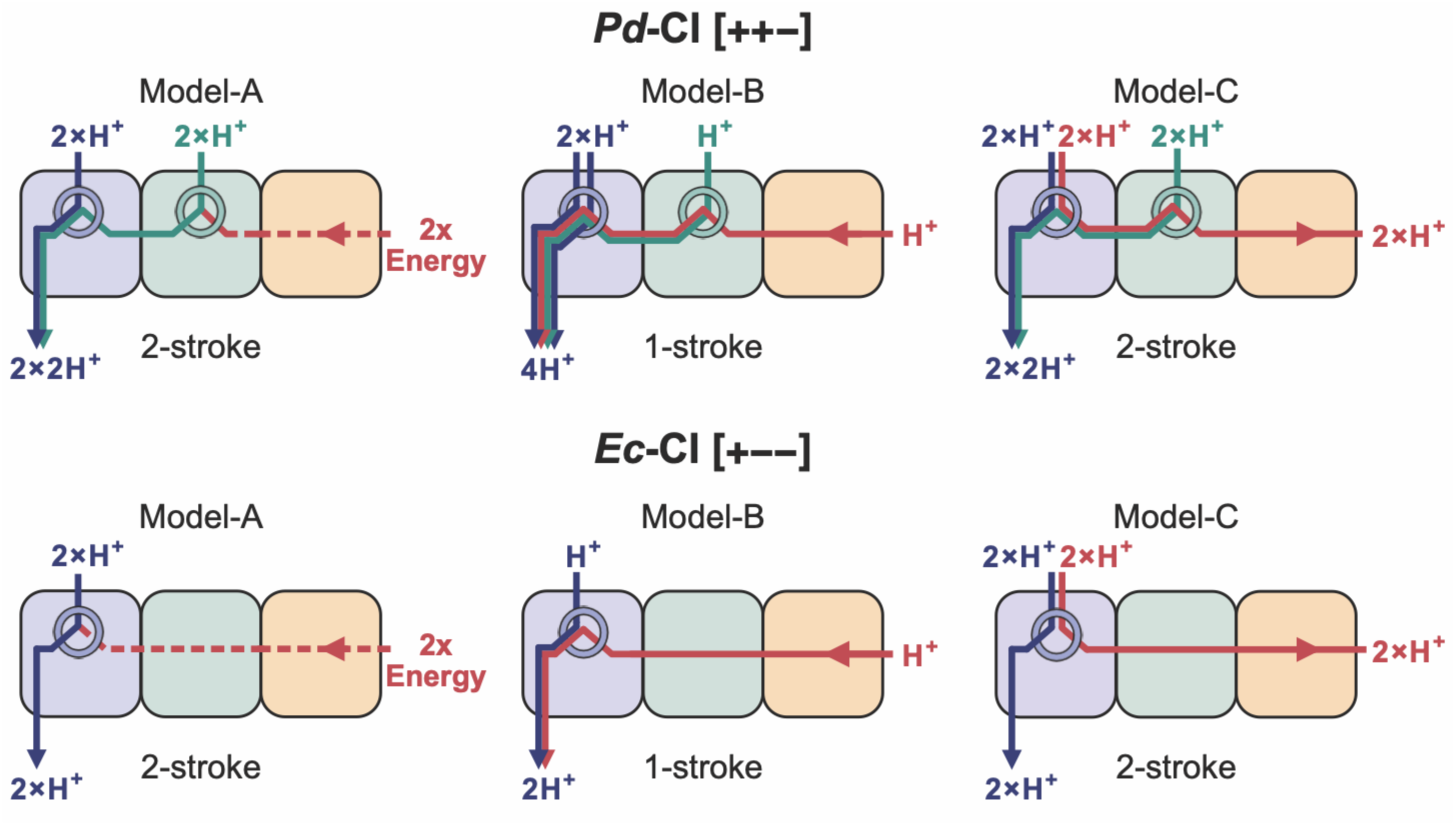
Three models for the modular mechanism of proton pumping by *P. denitrificans* [++-]CI and *E. coli* [+--]CI. In model-A, energy is transferred directly from redox catalysis, in model-B a high-energy proton is transferred from redox catalysis, and in model-C redox catalysis abstracts protons from the proton-pumping domain. Model-A and model-C are two-stroke mechanisms (the same proton-pumping cycle repeats twice for each NADH oxidized) whereas model-B is a one-stroke mechanism. Proton pumping is gated at two His-junctions (circled) in ND5 and ND4 in *Pd*-CI, but only in ND5 in *Ec*-CI. ND5 is in blue, ND4 in green and ND2 in orange, with the ND5-uptake protons in blue, the ND4-uptake protons in green, and energy-linked protons / energy transfer in red. The four-proton [++-]CI exemplified by *Pd*-CI is in the top row and the two-proton [+--]CI exemplified by *Ec*-CI is in the bottom row; the modular mechanism of proton translocation must be adaptable to both.

## Materials and Methods

### Mutagenesis in the *P. denitrificans* genome

Mutations were created in the genome of *P. denitrificans* by adapting a published protocol (*18*, *22*, *23*). Insertion cassettes were designed with 1000 base-pair homologous flanking regions upstream and downstream of *nqo12* (ND5), *nqo13* (ND4) or *nqo14* (ND2) and a kanamycin (kanR) selection marker before the upstream flanking region. Mutations were generated in the cassettes and inserted into a lacZ-containing pRVS1 backbone by Gibson assembly. The resulting plasmids were transformed into MFDpir, an *E. coli* donor strain, and conjugated into a *Pd*1222–ΔHy–Nqo5His6–pLMB509-NDH-2 parental strain (*22*) at a donor-recipient ratio of 1:10. *P. denitrificans* cells were grown with NDH-2 expression induced by 10 mM taurine throughout. The first recombination event following conjugation was selected for on Luria–Bertani (LB)-taurine agar plates containing 50 µg mL^-1^ kanamycin, and integration of the mutated gene checked by Sanger sequencing. Positive colonies were then grown overnight in LB-taurine without kanamycin and plated onto LB-taurine agar plates containing 200 µg mL^-1^ X-gal. Successful second recombination events, observed as white colonies, were screened by PCR amplification and sequencing across the mutation site. Piericidin A (5 µM) was included in media for overnight growth when second recombination events were repeatedly unsuccessful. Finally, the CI operon was checked for off-target mutations, using colony PCR amplification of the operon in two fragments followed by gel electrophoresis and Nanopore sequencing (Plasmidsaurus Inc.). The cell growth characteristics of all confirmed variants were checked by monitoring cell density via the 600 nm absorbance in a BMG CLARIOstar plus microplate reader. 1 mL aliquots of overnight pre-cultures were pelleted (16,000 x g, 1 min) and resuspended in LB without antibiotics or taurine to OD_600_ = 0.2. Then, 100 µL of suspension was added to 100 µL LB with or without 10 mM taurine, to give a starting OD_600_ of 0.1. Plates were incubated at 30 °C with 500 rpm orbital shaking and readings every 10 min and growth curves fitted using linear regression on logarithmically plotted data in QurvE v1.1 (*54*). Typical doubling times were 160 – 200 min and 200 – 240 min for catalytically active and inactive *Pd*-CI variants, respectively, with maximum OD_600_ of 1.8 – 2.2. Values outside these ranges were observed for six variants, which we then aimed to recreate. Three catalytically inactive variants, H263S^ND5^ (348 ± 41 min, OD_600_ 1.65 ± 0.08), K217Q^ND2^ (329 ± 16 min, OD_600_ 2.15 ± 0.10) and F354A:K310Q^ND4^ (254 ± 6 min, OD_600_ 2.16 ± 0.10) were not successfully recreated. However, three catalytically active variants that were recreated (H246Y^ND4^, F354A^ND4^ and T239A^ND2^) then displayed normal growth characteristics and the properties of the CI were not changed, consistent with secondary effects on cell growth originating outside the CI operon.

### Membrane preparations from *P. denitrificans*

*P. denitrificans* membranes were prepared based on a protocol described previously (*23*). 50 µL of an overnight preculture were used to inoculate 50 mL LB with 10 mM taurine in a 250 mL conical flask. Each flask was incubated at 30 °C with shaking at 225 rpm until late-log phase (OD_600_ 3.0 – 4.5). All subsequent steps were performed at 4 °C. Cells were harvested by centrifugation (10,000 x g, 7 min) and resuspended in 1 mL buffer containing 50 mM 2-(N-morpholino)ethanesulfonic acid (MES) (pH 6.5 at 4 °C), 0.002% w/v phenylmethylsulphonyl fluoride (PMSF) and the cOmplete™ EDTA-free protease inhibitor cocktail (Roche). Cells were lysed using a Q700 Qsonica probe sonicator with a 1.6 mm microtip for a total running time of 1 min (cycles of 5 s on and 45 s off) at 60% amplitude. Cell debris were removed (16,000 x g, 15 min) before membranes were collected by centrifugation (220,000 x g, 1 hour), then resuspended and homogenized in 50 mM MES (pH 6.5 at 4 °C). All mutant *P. denitrificans* strains were grown and membranes prepared alongside WT controls.

### Preparation of SBPs from *P. denitrificans*

*P. denitrificans* sub-bacterial particles (SPBs) were prepared as described previously (*18*, *23*). 500 µL of an overnight preculture were used to inoculate 1 L of succinate minimal media with 10 mM taurine in a 2 L conical flask. The media (adjusted to pH 7.2) contained 50 mM succinate, 9.35 mM NH_4_Cl, 2 mM MgSO_4_, 0.07 mM CaCl_2_, 0.29 mM KH_2_PO_4_, 0.69 mM K_2_HPO_4_, 25.2 mM Na-HEPES, 19.6 μM Na_2_-EDTA, 9 μM FeSO_4_, 0.1 μM MnCl_2_, 0.8 μM CuCl_2_, 1 μM Na_2_MoO_4_, and 2.5 μM ZnCl_2_. Each flask was incubated at 30 °C with shaking at 225 rpm to an OD_600_ of 1.5 – 2.5. All subsequent steps were performed at 4 °C. Cells were collected by centrifugation (13,690 x g, 10 min) then resuspended in ∼400 mL of buffer containing 10 mM Tris-SO_4_ (pH 7.5 at 4 °C) and 150 mM NaCl and recentrifuged (13,690 x g, 15 min). The pellet was resuspended to OD_600_ = 7.5 in ∼250 mL of buffer containing 10 mM Tris-SO_4_ (pH 7.5 at 4 °C) and 500 mM sucrose, then lysozyme (Merck Life Science Ltd.) was added to 250 µg mL^-1^ and the suspension incubated on ice for 1 hour. The treated cells were collected again by centrifugation (13,690 x g, 10 min) and resuspended in 50 mL of buffer containing 10 mM Tris-SO_4_ (pH 7.5 at 4 °C) and 500 μM MgSO_4_, then further buffer was added to a total volume of 400 mL, causing cell rupture and membrane inversion to form SBPs. After 15 min on ice, MgSO_4_ to 5 mM and a few flakes of DNase were added and the mixture incubated for a further 15 min. Remaining intact cells were removed (13,690 x g, 2 x 15 min) and the SBPs collected by centrifugation (13,690 x g, 1 hour). The SBP pellet was resuspended in 500 µL of buffer containing 10 mM Tris-SO_4_ (pH 7.5 at 32 °C) and 250 mM sucrose, aliquoted and flash frozen in liquid nitrogen for storage at −70 °C.

### Purification of complex I from *P. denitrificans*

*Pd*-CI was purified based on a protocol described previously (*14*, *22*). 2 L flasks containing 1 L of LB with 10 mM taurine were inoculated with 1 mL of an overnight preculture and grown at 30 °C with shaking at 225 rpm until late-log phase (OD_600_ 3.0 – 4.5). All subsequent steps were at 4 °C. Cells from 4–6 L of media were pelleted by centrifugation (6000 x g, 30 min) and resuspended in ∼400 mL of buffer containing 50 mM MES (pH 6.5 at 4 °C), 0.002% w/v PMSF and the cOmplete™ EDTA-free protease inhibitor cocktail. The cells were homogenized and lysed in a CF2 4 kW cell disruptor (Constant Systems Ltd) at 15,000 psi then 30,000 psi twice. Unlyzed cells were pelleted (29,750 x g, 1 hour) and the membranes collected by centrifugation (234,800 x g, 2 hours) and resuspended in a minimal volume of 50 mM MES (pH 6.5 at 4 °C). The solution composition was then adjusted to 20 mM MES (pH 6.5 at 4 °C), 150 mM NaCl, 2.5 mM CaCl_2_, 10 % glycerol, 0.002% PMSF and cOmpleteTM EDTA-free protease inhibitor cocktail (Roche) at a protein concentration of 9.5 mg mL^-1^ and the membranes solubilized for 30 min on ice by addition of 2.85% n-dodecyl-β-D-maltoside (DDM, Anatrace). Insoluble material was removed by centrifugation (235,000 x g, 45 min), imidazole added to 20 mM and the solution filtered (0.45 µm) and loaded onto a HisTrap HP column (Cytiva) pre-equilibrated with buffer A [20 mM MES (pH 6.5 at 4 °C), 150 mM NaCl, 2.5 mM CaCl_2_, 10 % glycerol, 0.02% asolectin, 0.02% 3-[(3-cholamidopropyl)dimethylammonio]-1-propane-sulfonate (CHAPS), 0.1% DDM] plus 80 mM imidazole. The column was washed with buffer A plus 80 mM imidazole then eluted with buffer A plus 300 mM imidazole. The eluate was concentrated (Amicon Ultra-15, 100 kDa MWCO) to < 1 mL, loaded onto a pre-equilibrated Superdex 200 increase 10/300 GL column (Cytiva) and eluted in buffer C [20 mM MES (pH 6.5 at 4 °C), 150 mM NaCl, 2.5 mM CaCl_2_, 10 % glycerol, 0.05% DDM]. CI peak fractions were collected, concentrated to 10-25 mg mL^-1^ (Amicon Ultra-0.5, 100 kDa MWCO) and flash frozen with 20% glycerol added for storage.

### Mutagenesis in the *nuo* operon on an *E. coli* expression plasmid

The *nuo* operon was deleted from the genome of *E. coli* strain BW25113 by allelic exchange (*55*) with a deletion cassette containing ∼850 base-pair homology regions upstream and downstream of the deletion. The cassette was assembled into a pFOK backbone by Gibson assembly and propagated in *E. coli* strain EC100D (+) *pir-116*. The resulting plasmid was transformed into the *E. coli* JKe201 donor strain, which was then conjugated with the parental BW25113 *E. coli* strain. Recombination of the pFOK plasmid into the genome of BW25113 was selected for on LB-agar plates containing 50 µg mL^-1^ kanamycin. Second recombination events (the elimination of the pFOK backbone) were selected by plating cells onto LB-agar with 10% w/v sucrose (the substrate for SacB), no sodium chloride, and 0.5 µg mL^-1^ anhydrotetracycline (to promote dual-negative selection by SacB and the endonuclease I-SceI). Colony PCR was used to confirm the deletion. The *nuo* operon was deleted in two stages, first generating a Δ*nuoA-G* strain then deleting *nuoH-N* to generate the BW25113-Δ*nuo* strain. Mutations were generated in subunits L (ND5), M (ND4) and N (ND2) in a pMA vector containing the *nuoLMN* fragment using the QuikChange II site-Directed Mutagenesis Kit (Agilent). Each mutated *nuoLMN* fragment was then inserted into a pBAD30 backbone containing *nuoA-K* by Gibson assembly, propagated in the NEB^®^ 10-beta *E. coli* strain, and transformed into the BW25113-Δnuo strain for expression.

### Membrane preparations from *E. coli*

Over-night precultures of *E. coli* cells were grown in 5 mL of LB containing 100 µg mL^−1^ of ampicillin at 30 °C, with shaking at 225 rpm. 500 µL of each preculture was then used to inoculate 50 mL of LB containing 100 µg mL^−1^ ampicillin in a 250 mL conical flask and the cultures grown at 30 °C with shaking at 225 rpm until early-log phase (OD_600_ 0.5 to 0.8, typically 4 hours), when expression of the *nuo* operon was induced by addition of 0.002% w/v L-arabinose. Cell growth was maintained under the same conditions until late-log phase (OD_600_ 1.5 to 2.5, typically 2 hours). Cell harvesting, lysis and membrane preparation were then carried out using the methods described above for *P. denitrificans*, with membranes resuspended in 50 mM MES (pH 6.5 at 4 °C).

### Preparation of SBPs from *E. coli*

For comparison of WT and variants of *Ec*-CI (Figure 4D and Supplementary Figure 13D), *E. coli* cells were grown overnight with shaking at 225 rpm at 30 °C, by using 2 x 5 mL of preculture to inoculate 2 x 500 mL of LB containing 100 µg mL^−1^ ampicillin and 0.002% w/v L-arabinose in 2 L flasks. For stoichiometry measurements (Figure 4D and Supplementary Figure 14), the parental *E. coli* strain was grown with shaking at 225 rpm at 30 °C until late exponential phase (OD_600_ ∼1.5), by using 2 x 5 mL of preculture to inoculate 2 x 500 mL of LB in 2 L flasks. All subsequent steps were at 4 °C. Cells were harvested by centrifugation (4,000 x g, 20 min) then resuspended in 20 mL (per 1 L of culture) of buffer containing 50 mM MES (pH 6.5 at 4 °C), 5 mM MgCl_2_ and 10% (v/v) glycerol. A few flakes of DNase (Roche) and a cOmplete™ EDTA-free protease inhibitor tablet (Roche) were added, and the suspension passed twice through a pre-chilled SPCH-10 pressure cell homogenizer (Stansted) at 12,000 psi. The resulting lysate was centrifuged (16,000 x g, 30 min) to remove cell debris, then again (100,000 x g, 1 hour) to collect the SBPs. The SBPs were resuspended in 1 mL of the same buffer using a glass Dounce homogenizer, then aliquoted and flash frozen in liquid nitrogen for storage at −70 °C.

### Preparations for coupled-enzyme assays

Fumarate hydratase (FumC (*56*)) and oxaloacetate decarboxylating malic enzyme (MaeB (*57*)) for monitoring succinate oxidation were prepared as described previously (*58*) in 10 mM Tris-SO_4_ (pH 7.0 at 4 °C), 5 mM EDTA and 1 mM DTT, and 60 mM Tris-SO_4_ (pH 7.4 at 4 °C), 20 mM β-mercaptoethanol and 10% (w/v) glycerol, respectively. The alternative quinol oxidase from *Trypanosoma brucei brucei* (AOX) was expressed recombinantly in *E. coli* and purified as described previously (*59*, *60*) in 20 mM Tris-HCl (pH 8.0 at 4 °C), 50 mM MgSO_4_, 160 mM NaCl, 20% (v/v) glycerol and 0.042% DDM. Methods for producing the cytochrome *bo*_3_ oxidase from *E. coli* (*bo*_3_) were based on published protocols (*61*). *bo*_3_ was expressed from a pET-17b plasmid encoding the cyoABCDE operon in a ΔcyoBCD C43(DE3) strain of *E. coli*, with a His_6_-tag at the C-terminus of subunit II, generously provided by Prof. Christoph von Ballmoos (Universität Bern). 6 x 1 L of media containing 2 g L^-1^ (NH_4_)_2_SO_4_ and 13.6 g L^-1^ K_2_HPO_4_, (adjusted to pH 7), plus 5 g L^-1^ FeSO_4_, 1 mM MgSO_4_, 10 µM CuSO_4_, 0.2% NH_4_Cl, 0.2% glucose, 100 mg L^-1^ thiamine, 100 mg L^-1^ ampicillin in 2 L flasks were inoculated using an overnight pre-culture in LB and cells grown at 37 °C and 225 rpm shaking. Expression was induced at exponential phase (OD_600_ 0.4–0.6) with 0.5 mM IPTG, followed by 12 hours growth at 30 °C. Cells were harvested by centrifugation (5,000 x g, 20 min) and resuspended in membrane buffer (50 mM K_2_HPO_4_ (pH 8.3 at 4 °C), 0.002% PMSF (w/v) and the Roche EDTA-free cOmplete protease inhibitor cocktail). All subsequent steps were performed at 4 °C. The cells were passed through a Z-plus 2.2 kW cell disruptor (Constant Systems Limited) once at 15,000 psi and twice at 30,000 psi. Debris were removed by centrifugation (20,000 x g, 1 hour) then the membranes collected (225,000 x g, 2 hour), resuspended in membrane buffer to 30 mg mL^-1^ and flash frozen for storage at -70 °C. Thawed membranes were diluted to 10 mg mL^-1^, with the buffer composition corrected to 50 mM K_2_HPO_4_ (pH 8.3 at 4 °C), 150 mM NaCl, 0.002% PMSF (w/v), and 2% DDM (w/v), and solubilized for 1 hour with constant stirring. Insoluble material was removed by centrifugation (250,000 x g, 30 min) and the supernatant filtered (0.22 μm syringe filter MillexHP, Millipore) and loaded onto a 5 mL His-TrapTM HP column (Cytiva) equilibrated in buffer containing 25 mM K_2_HPO_4_ (pH 8.3 at 4 °C), 150 mM NaCl and 0.1% DDM (w/v). The column was washed with 50 mL buffer, then eluted over a 100 mL gradient from 0 mM to 300 mM imidazole. The fraction eluting at 150 mM imidazole was concentrated with a 50 kDa MW cut-off centrifugal filter (Merck Millipore) and exchanged into 25 mM K_2_HPO4 (pH 8.3 at 4 °C), 0.1% DDM (w/v) using a PD-10 desalting column (Cytvia) before being loaded onto a 5 mL HiTrap™ Q HP anion exchange column (Cytiva). Following a 50 mL gradient from 0 to 1 M NaCl, the material eluting at 200–250 mM NaCl was pooled, concentrated, and loaded on a Superdex 200 Increase 10/300 GL size exclusion column (Cytiva), equilibrated with 25 mM K_2_HPO_4_ (pH 8.3 at 4 °C), 150 mM NaCl and 0.03% DDM (w/v). The peak fraction was concentrated to ∼10 mg mL^-1^ and glycerol added to 30% (v/v) before flash-freezing in liquid nitrogen. Proteoliposomes containing *bo*_3_ (*bo*_3_-PLs) or AOX (AOX-PLs) were prepared using a protocol adapted from Grba et al. (*31*). A 25 mg mL^-1^ lipid chloroform solution was created by mixing 1,2-dioleoyl-sn-glycero-3-phosphocholine (DOPC, Avanti Polar Lipids), 18:1 cardiolipin (CDL, Avanti Polar Lipids) and 1,2-dioleoyl-sn-glycero-3-phosphoethanolamine (DOPE, Avanti Polar Lipids) in an 8:1:1 weight ratio. 10 mg of lipids were dried under N_2_ in a glass tube then rehydrated in 1 mL of buffer containing 10 mM MOPS (3-(N-morpholino)propanesulfonic acid, pH 7.5, 20 °C) and 50 mM KCl. Liposomes were formed using a Q700 Qsonica probe sonicator with a 1.6 mm microtip (total 2.5 min with 15 s on and 30 s off cycles at 60% amplitude) then partially solubilized in 0.5% (w/v) sodium cholate. Following 10 min incubation on ice, 200 μg of *bo*_3_ or 600 μg of AOX were added and incubated for a further 10 min. The *bo*_3_/AOX-PLs were exchanged back into 10 mM MOPS (pH 7.5, 20 °C) and 50 mM KCl using a pre-equilibrated PD-10 desalting column (Cytiva) to remove detergents, centrifuged (210,000 x g, 1 hour, 4 °C) and resuspended in 100 µL, then their concentrations determined using the BCA assay before flash freezing and storage.

### Catalytic activity assays

All catalytic activity assays were performed at 32 °C in a SpectraMax ABS plus plate reader (Molecular Devices). *P. denitrificans* membranes, *Pd*-CI and *E. coli* membranes were assayed in buffer containing 10 mM MES (pH 6.5 at 32 °C), 25 mM NaCl and 2 mM CaCl_2_. *P. denitrificans* SBPs were assayed in 10 mM Tris-SO_4_ (pH 7.5 at 32 °C) and 250 mM sucrose with 0.5 µg mL^-1^ gramicidin (Sigma-Aldrich) when required to dissipate Δp. The assays on *E. coli* SBPs reported in Supplementary Figure 13D were in 10 mM MOPS (pH 7.5 at 25 °C) and 50 mM KCl with 2 µM FCCP (carbonyl cyanide 4-(trifluoromethoxy)phenylhydrazone, Sigma-Aldrich) to dissipate Δp. 15–20 µg mL^-1^ alamethicin (Cayman Chemicals) was added to membrane assays to ensure substrate access to all CI molecules. 0.15% w/v asolectin (Avanti Polar Lipids) and 0.15% w/v CHAPS were included in *Pd*-CI assays to maintain enzyme solubility. dNADH was either purchased (Sigma-Aldrich) or produced in-house by enzymatic reduction of dNAD^+^ by alcohol dehydrogenase from *Saccharomyces cerevisiae* (Sigma Aldrich), by adapting a protocol described previously (*62*). Rates of dNADH:APAD^+^ oxidoreduction (APAD^+^ is 3-acetylpyridine adenine dinucleotide, an NAD^+^ analogue) were monitored at 400–450 nm (ε = 3.16 mM^-1^ cm^-1^) with 100 µM dNADH, 500 µM APAD^+^ (Sigma-Aldrich) and piericidin A (1 µM for *P. denitrificans* or 2 µM for *E. coli*) to prevent CI-mediated quinone reduction, with either 10 µg mL^-1^ membranes or SPBs or 0.5 µg mL^-1^ CI. Rates of dNADH:O_2_ oxidoreduction by membranes and SBPs were monitored at 340–380 nm (ε = 4.81 mM^-1^ cm^-1^) with 100 µM dNADH and 5 (*P. denitrificans*) or 10 (*E. coli*) µg mL^-1^ membranes or SBPs and controlled by addition of 5 µM piericidin A to inhibit CI. Rates of dNADH:decylubiquinone (DQ) oxidoreduction by purified *Pd*-CI were monitored at 340–380 nm (ε = 4.81 mM^-1^ cm^-1^) with 100 µM dNADH, 100 µM DQ and 0.5 µg mL^-1^ CI, and controlled by addition of 5 µM piericidin A. Rates of RET (succinate:dNAD^+^ oxidoreduction) by *Pd*-SBPs were monitored at 340–380 nm (ε = 4.81 mM^-1^ cm^-1^) in buffer containing 10 mM Tris-SO_4_ (pH 7.5 at 32 °C), 250 mM sucrose, 2 mM MgSO_4_ and 1 mM K_2_SO_4_ with 3 mM dNAD^+^, 5 mM succinate and 20 µg mL^-1^ SBPs and controlled by addition of 2 µM piericidin A, 1.25 µM myxothiazole and 400 µM KCN, or 0.5 μg mL^−1^ gramicidin. For sets of biological replicates (N ≥ 3), statistical differences to the *Pd*-WT or *Ec*-WT were calculated by one-way Welch’s ANOVA using Dunnett’s T3 test against all *Pd*-or *Ec*-variants (* p < 0.05, ** p < 0.01 to **** p < 0.0001, otherwise non-significant). Statistical analyses are not reported for sets of technical replicates (n ≥ 3), when the comparison requires data from only one biological replicate or preparation for each variant.

### ACMA fluorescence quenching (see also Supplementary Figure 5)

Proton pumping by *Pd*-CI and *Ec*-CI variants was assessed by the quenching of 9-amino-6-chloro-2-methoxyacridine (ACMA) fluorescence in energized SBPs using a CLARIOstarPlus plate reader (BMG Labtech) at 25 °C. For *Pd*-CI, four respiratory pathways were compared: CI-CIII-CIV (dNADH:O_2_), CI-AOX (dNADH:O_2_), NDH2-CIII-CIV (NADH:O_2_) and NDH2-AOX (NADH:O_2_). ACMA fluorescence was monitored at λ_ex_ = 419-15 nm and λ_em_ = 484-20 nm, using black-bottomed 96-well plates in 10 mM MOPS (pH 7.5 at 25 °C) and 50 mM KCl with 0.5 µM ACMA and 0.1 µM valinomycin. SBPs were added at 5 µg mL^−1^ and, where present, AOX was added at 2.5 µg mL^−1^ along with 1.25 µM myxothiazol and 400 µM KCN to inhibit CIII-CIV. For NDH2-dependent pathways CI was inhibited by 2 µM piericidin A, and for CI-dependent pathways dNADH was used instead of NADH. For *Ec*-CI, only two dNADH:O_2_ pathways were assessed: CI-oxidases (referring to the native oxidases present) and CI-AOX. *E. coli* SBPs were added at 20 µg mL^−1^ and AOX at 10 µg mL^−1^, with native oxidase activities inhibited by 10 mM KCN. Following a 5-min pre-incubation, catalysis was initiated by injection of 50 µM (*Pd*-CI) or 100 µM (*Ec*-CI) dNADH or NADH (from a 1 mM stock solution) and 1 s orbital mixing, the fluorescence followed for 180 s, then 50 mM NH_4_Cl (from a 1M stock solution) was injected and mixed for 2 s, and the fluorescence followed for a further 60 s. Traces were normalized (F/F_0_) to the average fluorescent value over the first 60 seconds prior to substrate addition. At least three technical replicates were recorded for each pathway. Corresponding NADH oxidation activities were determined under identical conditions and monitored at 340 nm (ε = 6.22 mM^−1^ cm^−1^) to minimize interference from the ACMA.

### Stoichiometry measurements using SBPs from *P. denitrificans* (see also Supplementary Figure 8)

Measurements were based on protocols described previously (*18*, *23*, *63*) and carried out at 32 °C using 20-30 µg mL^-1^ SBPs in buffer containing 10 mM Tris-SO_4_ (pH 7.5, 32 °C), 250 mM sucrose, 2 mM MgSO_4_, 1 mM K_2_SO_4_, 10 mM K_2_PO_4_, 1 mM ADP (Sigma-Aldrich) and 100 µM Ap5A [P1, P5-di(adenosine-5′) pentaphosphate (Sigma-Aldrich), to inhibit adenylate kinase]. ADP was pre-treated to remove ATP by incremental addition of glucose to 50 mM ADP in 20 mM HEPES (pH 7.8 at 20 °C), 20 mM KCl and 5 mM MgCl_2_ with 2 U mL^-1^ hexokinase, followed by heat-inactivation of the hexokinase (80°C, 15 min). Rates of dNADH:O_2_ oxidoreduction by CI-CIII-CIV were monitored at 340–380 nm (ε = 4.81 mM^-1^ cm^-1^) upon addition of 200 µM dNADH and modulated using piericidin A. Rates of succinate:O_2_ oxidoreduction were monitored using a coupled assay system (*58*) in which succinate oxidation (5 mM) was coupled to NADP^+^ reduction (2 mM, monitored at 340–380 nm, ε = 4.81 mM^−1^ cm^−1^) by 180 µg mL^-1^ FumC and 0.9 mg mL^-1^ MaeB and modulated using atpenin A5. 2 µM piericidin A was added to inhibit *Pd*-CI catalysis, and the linear rates were fitted after 3 min to eliminate any lag phase from the coupled enzymes. Rates of dNADH:O_2_ oxidoreduction by CI-*bo*_3_ were catalyzed by the *Pd*-CI in SBPs (with CIII-CIV inhibited by 1.25 µM myxothiazol and 100 µM KCN) and 2.5 µg mL^-1^ *bo*_3_-PLs, mediated by 10 µM ubiquinone-1 (Sigma-Aldrich), with the rate modulated using piericidin A. Rates of ATP synthesis were measured by withdrawing 10 µL aliquots of assay mixtures at 30 s intervals for 3 minutes, starting from 1 (dNADH) or 3 min (succinate, to avoid any lag-phase in the coupled enzymes for succinate detection) after catalysis was initiated. The aliquots were quenched by adding them to 40 µL of 4% (v/v) trifluoroacetic acid, which was neutralized 20 s later by addition of 250 µL of 1M Tris-SO_4_ (pH 8, 32 °C). Each neutralized solution was divided into three wells of a white, flat-bottomed 96-well luminescence plate and combined, by direct injection in a BMG Labtech CLARIOstar plus microplate reader, with 80 µL ATP detection reagent [ATP Bioluminescence Assay Kit CLS II (Roche) diluted 4x relative to the manufacturer’s instructions, or an in-house mix (*64*) of 1.87 mM DTT, 0.1 mg mL^-1^ bovine serum albumin (Sigma-Aldrich), 0.48 µg mL^-1^ luciferase (Sigma-Aldrich) and 30 µM luciferin (Sigma-Aldrich)]. The plate was shaken for 5 s to mix the solutions, then the luminescence measured for 10 s at 500-660 nm. Luminescence values were converted to ATP concentrations using a set of identically treated ATP standards that contained a total of 1mM ADP + ATP, with ATP concentrations from 0–100 µM and the resulting rates of ATP synthesis plotted against the rates of substrate oxidation. The *Pd*-CI stoichiometry (*n*_CI_) was determined, as described previously (*18*, *23*, *63*) by minimizing the least-squared errors for global fits of all three datasets together for candidate values of *n*_CI_, assuming pathway stoichiometries of (*n*_CI_ + 6) for CI-CIII-CIV, 6 for CII-CIII-CIV and *n*_CI_ for CI-*bo*_3_ (PLs). The *R*^2^ values are reported for the best fit for each dataset to indicate the quality of the fit, along with the plots of least-squared error vs. *n*_CI_ to indicate the confidence in the best-fit value.

### Stoichiometry measurements using SBPs from *E. coli* (see also Supplementary Figure 14)

The protocol for *Ec*-SBPs was adapted from that described above for *Pd*-SBPs. All stoichiometry data are presented for the parental strain of *E. coli* due to the SBP preparations being better coupled than for the WT expression strain, and the cells were harvested in exponential phase to ensure expression of the cytochrome *bo*_3_ oxidase rather than the cytochrome *bd* oxidases (*65*). The buffer composition, temperature, and assay duration were optimized to ensure the activity of the *Ec*-SBPs remained stable over the experimental timescale. Measurements were carried out at 25 °C using 40 µg mL^-1^ SBPs in buffer containing 50 mM MOPS (pH 6.5, 25 °C), 5 mM MgCl_2_, 1 mM K_2_SO_4_, 10 mM K_2_PO_4_, 9.8% glycerol, 1 mM ADP and 100 µM Ap5A. The ADP was pre-treated as described above. Rates of dNADH:O_2_ oxidoreduction by CI-*bo*_3_ were monitored at 340–380 nm (ε = 4.81 mM^-1^ cm^-1^) upon addition of 200 µM dNADH and modulated using piericidin A. Rates of dNADH:O_2_ oxidoreduction by CI-AOX were catalyzed by the *Ec*-CI in SBPs (with native quinol oxidase activity inhibited by 15 mM KCN) and 5 µg mL^-1^ AOX-PLs, mediated by 40 µM ubiquinone-1 (UQ_1_), with the rate modulated using piericidin A. Only two pathways could be compared because rates of succinate oxidation were too low to provide accurate data for the CII-*bo*_3_ pathway, and because a sufficiently specific *Ec*-CI inhibitor could not be identified to inhibit *Ec*-CI but not NDH2, in order to utilize the NDH2-*bo*_3_ pathway instead (see Supplementary Figure 14). Rates of ATP synthesis were measured by withdrawing 10 µL aliquots of assay mixtures at 20 s intervals for 2 minutes, starting from 20 s after catalysis was initiated. The aliquots were quenched and the ATP concentrations determined as described above to plot the rates of ATP synthesis against the rates of substrate oxidation. The *Ec*-CI stoichiometry (*n*_CI_) was determined, as described previously (*18*, *23*, *63*), by minimizing the least-squared errors for global fits of both datasets together for candidate values of *n*_CI_, assuming *E. coli* pathway stoichiometries of (*n*_CI_ + 4) for CI-*bo*_3_ (*39*, *40*) and *n*_CI_ for CI-AOX (PLs). The *R*^2^ values are reported for the best fit for each dataset to indicate the quality of the fit, along with the plots of least-squared error vs. *n*_CI_ to indicate the confidence in the best-fit value.

### Biochemical analyses

For SDS PAGE, *Pd*-CI samples were incubated for 10 min at ∼20 °C in 0.125 M Tris-HCl (pH 6.8 at ∼20 °C), 20% (w/v) glycerol, 4% (w/v) SDS, 0.005% (w/v) bromophenol blue and 0.1 M dithiothreitol, loaded onto a 4– 12% Bolt^TM^ Bis-Tris Plus Mini Protein Gel, and run according to the manufacturer’s instructions. Bands were visualized using 0.25% w/v Coomassie blue R250 alongside Precision Plus ProteinTM KaleidoscopeTM prestained standards. To evaluate CI stability, nanoDSF (differential scanning fluorimetry) traces were recorded using a Prometheus NT.48 (NanoTemper Technologies) as described previously (*14*, *23*). *Pd*-CI was diluted to 0.2 mg mL^−1^ in 20 mM MES (pH 6.5 at 4 °C), 150 mM NaCl, 10 mM CaCl_2_, 10 % v/v glycerol, 0.05 % w/v DDM, loaded into four capillaries, and the fluorescence monitored at 330 to 350 nm (λ_ex_ = 280 nm) as the samples were heated from 20 to 90 °C).

### Structure analyses

Structure coordinates from the Protein Data Bank and cryoEM density maps were analyzed using Coot v1.1 (*66*), UCSF ChimeraX (*67*) and CASTpFold (*68*).

## Acknowledgements

We thank Professor Christoph von Ballmoos (Universität Bern, Switzerland) for the cytochrome *bo*_3_ expression plasmid, Valentin Datcu and Ketul Saharan (University of Cambridge) for *bo*_3_ preparation, and Emily Smith and Spencer Regan-Smith (University of Cambridge) for contributing to the preparation and initial evaluation of *Pd*-CI mutants.

## Author contributions

R.A.W. performed all *P. denitrificans* mutagenesis, prepared *P. denitrificans* complex I and membranes, determined *P. denitrificans* membrane activities, and contributed to preparing figures. H.L.M. prepared *P. denitrificans* SBPs, and performed all RET and stoichiometry measurements. M.M.C.-L. performed all *E. coli* mutagenesis, sample preparation and membrane activity measurements with support from J.H.S. J.J.W. performed ACMA analyses, supported activity and stoichiometry measurements, and led the preparation of the figures. R.A.W. and J.H. conceived the project. J.H. supervised the project, acquired the funding and led the preparation of the manuscript with input from all authors.

