## Supplementary Information for "A modular mechanism determines the proton-pumping stoichiometry of respiratory complex I"

**This file includes:**
Supplementary Figures 1-14
Supplementary Table 1

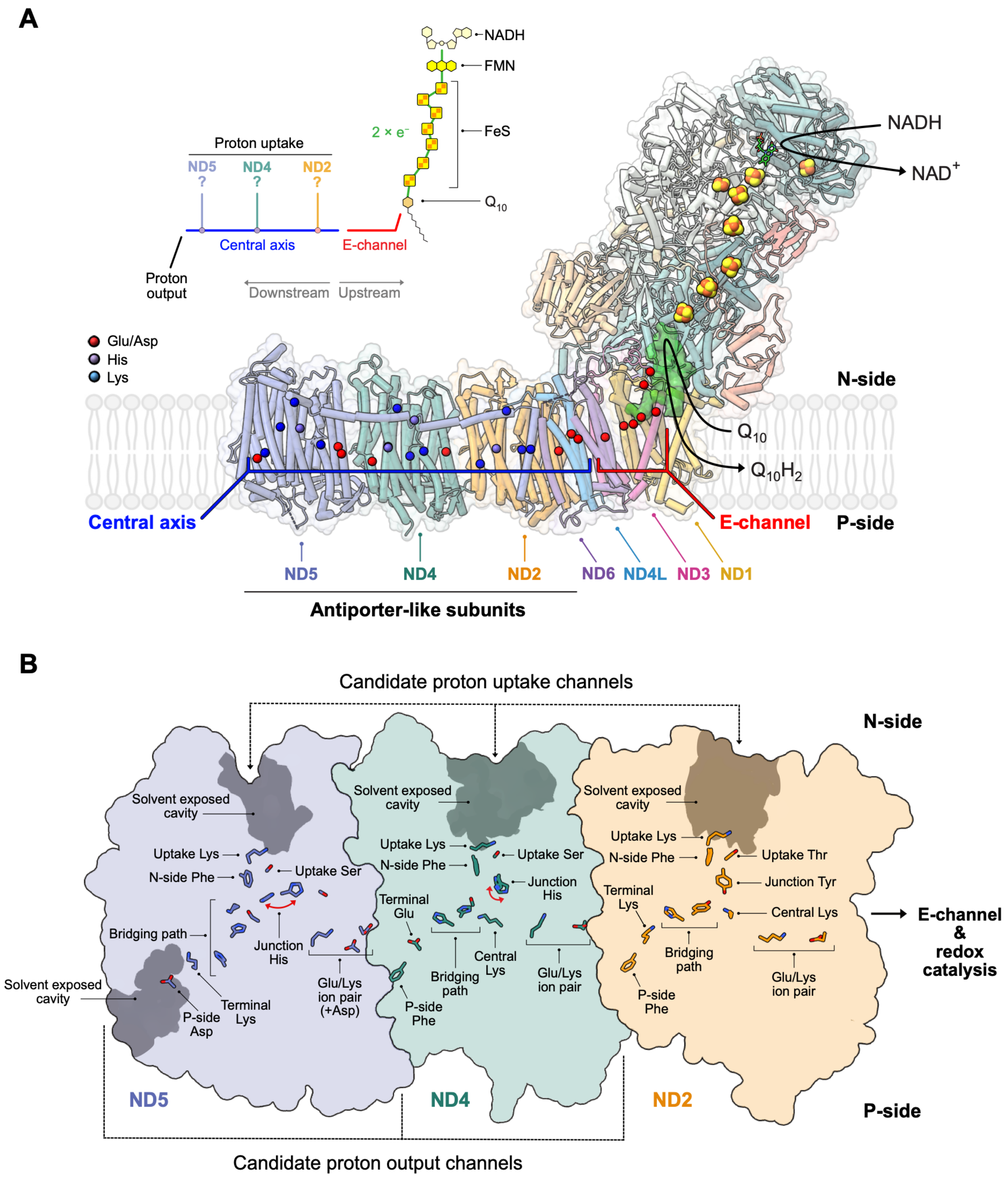


**Supplementary Figure 1. Summary of the structure and key functional elements and residues of complex I.** (**A**) The structure of complex I from *Paracoccus denitrificans* (*Pd*-CI, PDB: 8QBY) (*14*). NADH is oxidized at a flavin-containing active site at the top of the hydrophilic domain, and the electrons transferred by a chain of iron-sulfur clusters (orange and yellow) to the quinone-binding site (Q-site, green surface), where ubiquinone-10 ($Q_{10}$) is reduced. Residues in the membrane domain that form the E-channel and central axis and are crucial for coupling redox catalysis to proton pumping are shown. Note that we use the nomenclature for the subunits of human complex I (ND1-6 and ND4L) rather than the specific nomenclature for *P. denitrificans* (Nqo8, Nqo14, Nqo7, Nqo13, Nqo12, Nqo10 and Nqo11, respectively). Inset, simple summary of the major structurally defined elements and the 'downstream' direction of energy transfer during catalysis, from redox catalysis to the proton-pumping subunits. (**B**) The antiporter-like subunits (ND5, ND4, and ND2) showing the residues of the central axis and candidate proton-uptake/output pathways, and observed solvent-exposed cavities. Red arrows illustrate the movements of the junction-His in ND5 and ND4 observed in structures of mammalian complex I (*16*). This panel illustrates the relative positions of key functional elements and residues and establishes the nomenclature used for them.

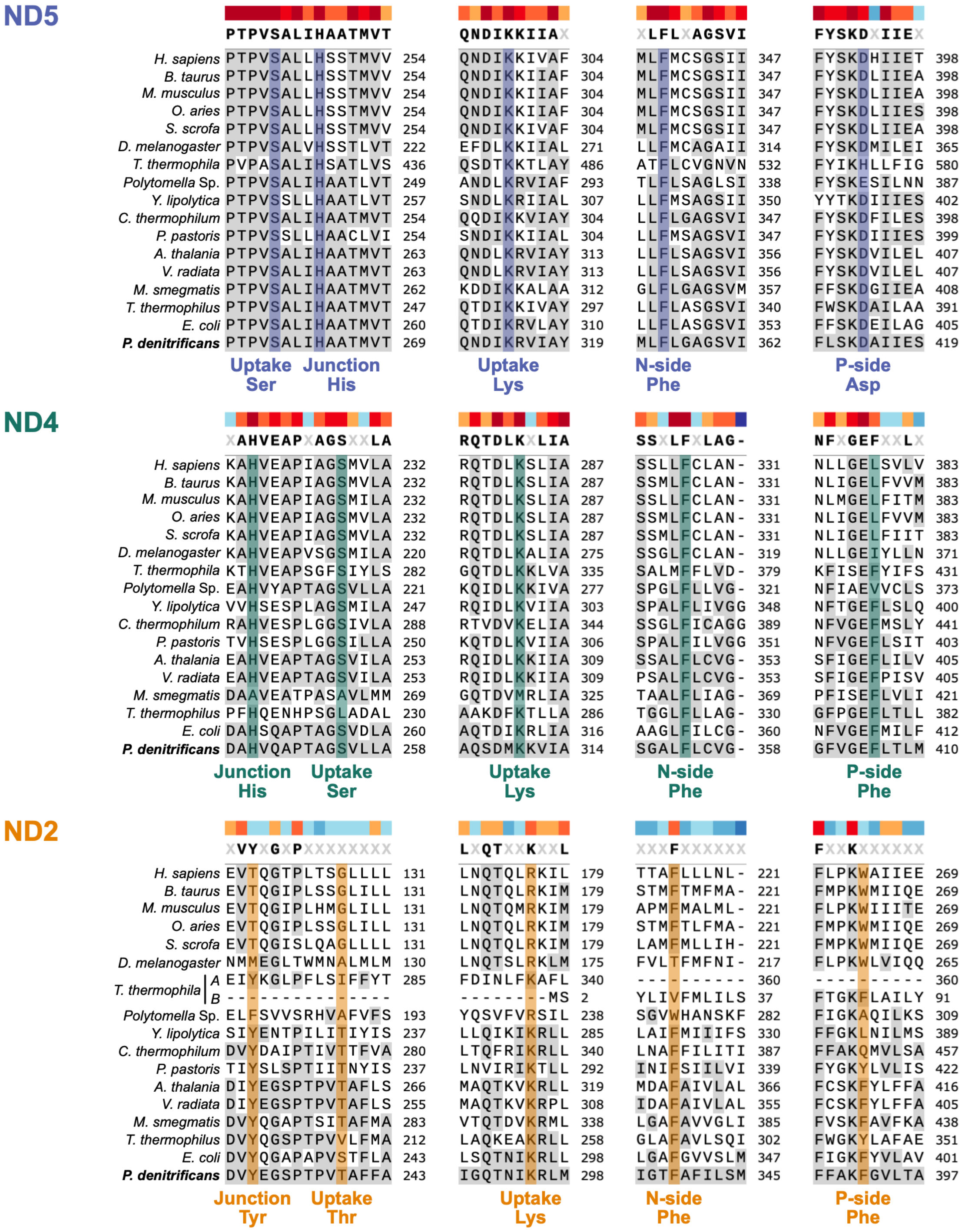

**Supplementary Figure 2. Sequence alignments for key residues in all species of complex I for which high-resolution structures (<3.5 Å) have been determined.** The consensus sequences (≥ 50% of the sequences shown) are given at the top with the level of conservation indicated by color (red, most highly conserved to blue, not conserved). Residues shaded in grey are conserved in *P. denitrificans* and residues of interest are highlighted and labelled. *T. thermophila* ND2 is split into two proteins (Ymf65/ND2A and ND2B) here labelled A and B for simplicity (*69*) and, due to poor local sequence alignment, the sequence of the transmembrane helix containing the N-side Phe$^{ND2}$ (W276) in *Polytomella* Sp. was manually realigned based on its structure. Sequences were aligned using Clustal Omega (*70*, *71*) and visualized in SnapGene. The P-side Asp$^{ND5}$, Phe$^{ND4}$ and Phe$^{ND2}$ are in structurally conserved positions in each subunit, with the P-side Asp$^{ND5}$ at the start of the structurally defined output pathway. Supplementary Table 1 gives the residue numbers in each species, along with full species names and structure references.

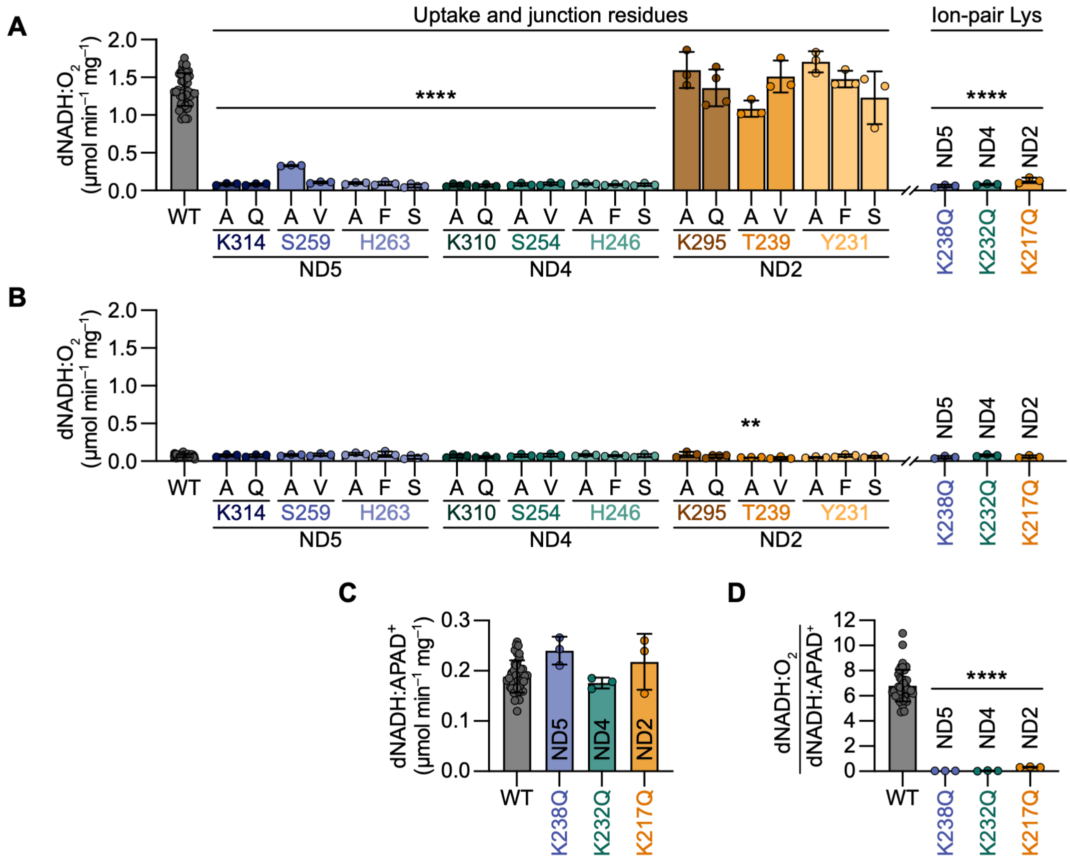


**Supplementary Figure 3. Supporting data for the catalytic activity measurements reported in Figure 1C, plus catalytic activity measurements on variants of the three ion-pair Lys in *P. denitrificans* CI.** Rates of dNADH:$O_2$ oxidoreduction without (**A**) and with (**B**) the CI inhibitor piericidin A (5 μM) were determined in independent *Pd*-membrane preparations ($N \geq 3$, ± S.D.). (**C**) Rates of dNADH:$APAD^+$ oxidoreduction determined in independent membrane preparations of three *Pd*-CI ion-pair Lys variants ($N \geq 3$, ± S.D.). (**D**) Normalized rates of *Pd*-CI turnover (panel A-B divided by panel C)) ($N \geq 3$, ± S.D.) for the *Pd*-CI ion-pair Lys variants. See Materials and Methods for experimental conditions and protocols.

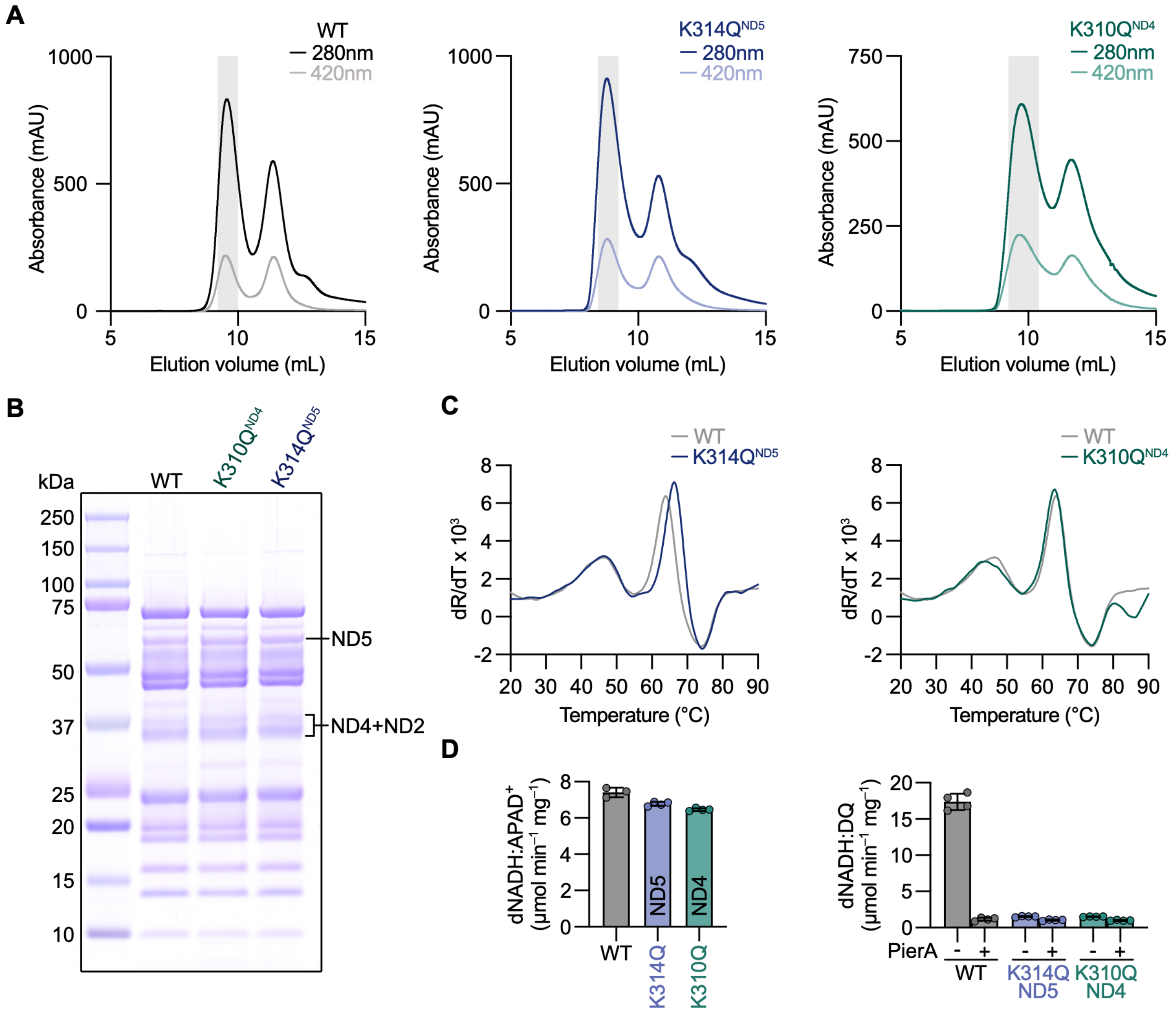


**Supplementary Figure 4. Isolation of intact, stable and homogeneous complexes for the catalytically inactive *Pd*-K314Q$^{ND5}$ and *Pd*-K310Q$^{ND4}$ variants.** (**A**) Size-exclusion chromatography traces recorded during the purification of the *Pd*-WT (black), *Pd*-K314Q$^{ND5}$ (blue) and *Pd*-K310Q$^{ND4}$ (green) complexes. Selected fractions are shaded. (**B**) SDS-PAGE analyses of the purified complexes with the bands for subunits ND5, 4 and 2 labelled. (**C**) NanoDSF traces reported as the first derivative of R with respect to temperature, where R is the ratio of the absorbance at 330 and 350 nm ($A_{330}/A_{350}$, average of four technical replicates) of the *Pd*-K314Q$^{ND5}$ (blue) and *Pd*-K310Q$^{ND4}$ (green) complexes compared to WT *Pd*-CI (grey). (**D**) Catalytic activity assays on the purified complexes: the dNADH:APAD$^+$ (left) and dNADH:decylubiquinone (dNADH:DQ, right) oxidoreductase activities in the presence and absence of 5 µM of the CI inhibitor piericidin A (PierA) ($n \geq 3$ technical replicates, ± S.D.). See Materials and Methods for experimental conditions and protocols.

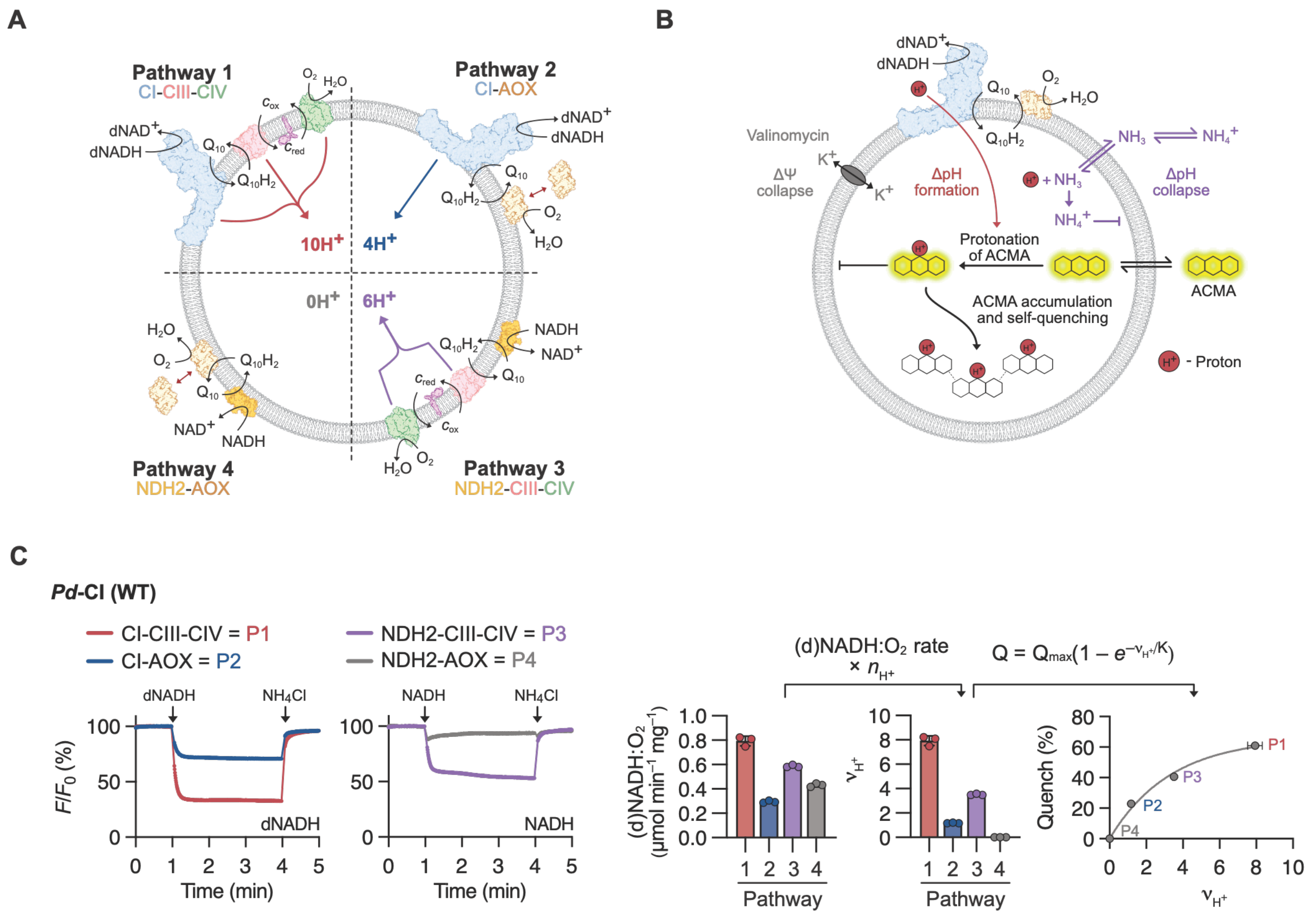


**Supplementary Figure 5. Principles of proton-pumping measurements in *Pd*-SBPs by ACMA fluorescence quenching.** (**A**) Four respiratory pathways used to analyze proton pumping in SBPs from *P. denitrificans*. Pathway 1 uses CI (4 $H^+$), CIII-CIV (6 $H^+$) to pump 10 $H^+$ per dNADH oxidized. Pathway 2 uses CI (4 $H^+$) and exogenous AOX (0 $H^+$, CIII and CIV are inhibited) to pump 4 $H^+$ per dNADH oxidized. Pathway 3 uses NDH2 (0 $H^+$, CI is inhibited), CIII-CIV (6 $H^+$) to pump 6 $H^+$ per NADH oxidized. Pathway 4 uses NDH2 and AOX (CI, CIII and CIV are inhibited) to pump 0 $H^+$ per NADH oxidized. (**B**) ACMA fluorescence quenching as a semi-quantitative measure of proton pumping. Valinomycin is added to dissipate ΔΨ so ΔpH builds to the total value of Δp. Unprotonated ACMA readily crosses the SBP membrane; during turnover it becomes protonated and trapped in the acidic vesicular lumen. Accumulation of ACMA in the lumen quenches its fluorescence. Addition of $NH_4Cl$ dissipates ΔpH through $NH_3$-mediated buffering of the lumen, allowing ACMA to escape and restoring its fluorescence. (**C**) ACMA traces for WT SBPs from *P. denitrificans* (average of three technical replicates, ± S.D.). Variable rates and extents of quenching are observed depending on the pathway and the magnitude of ΔpH. Rates of dNADH or NADH oxidation measured under matching conditions are converted to proton-pumping rates ($\nu_{H^+}$, µmol $min^{-1}$ $mg^{-1}$) by multiplying them by the stoichiometry of each pathway (assuming $n_{CI}$ = 4) (see bar charts). Finally, the extent of quenching (measured at 4 min) is plotted against the calculated proton-pumping rate and fit empirically using an exponential saturation equation $Q = Q_{max}(1 - e^{-\nu_{H^+}/K})$, where Q is the measured ACMA fluorescence quench (%), $Q_{max}$ is the maximum attainable quench, and $K$ is a constant that increases with proton leak (higher proton-pumping rates are needed to achieve the same quench). Because the ACMA signal saturates, Q is not linearly dependent on proton-pumping rate – but it always increases with it, and the fit shown for WT *Pd*-CI shows that the data are consistent with $n_{CI}$ = 4. When Q does not increase with the calculated proton-pumping rate, the data indicate that the value of $n_{CI}$ used is not correct (see *Pd*-H246Y$^{ND4}$ and *Pd*-K263Q$^{ND4}$ in Supplementary Figure 9 and *Pd*-F354A$^{ND4}$ in Supplementary Figure 11A). See Materials and Methods for experimental conditions and protocols.

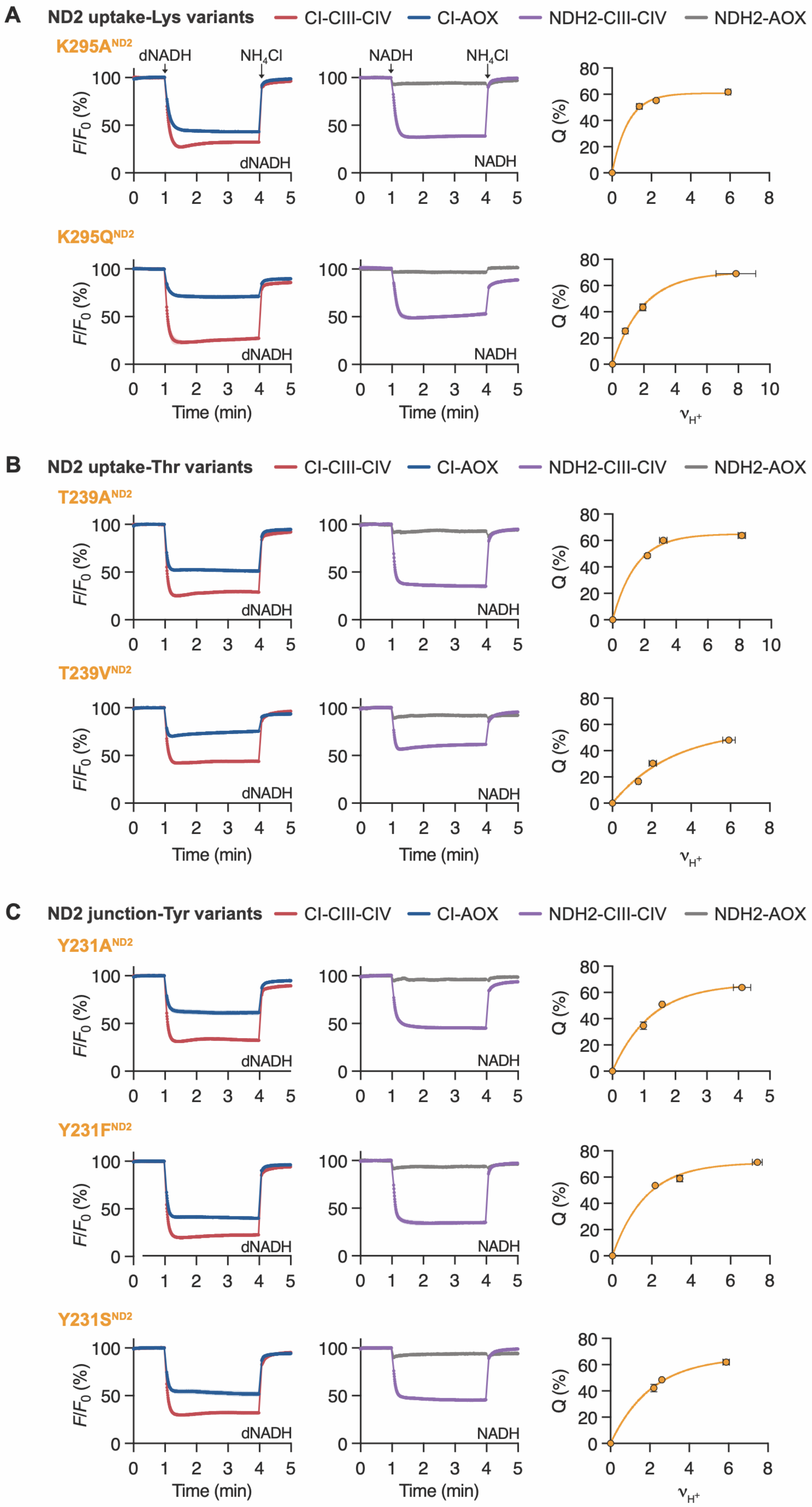


**Supplementary Figure 6. Proton pumping by the ND2-uptake pathway variants of *Pd*-CI in Figure 1 assessed by ACMA fluorescence quenching in *Pd*-SBPs.** ACMA quenching by the ND2-uptake pathway variants was analyzed as described in Supplementary Figure 5, and exemplified for WT *Pd*-CI in Supplementary Figure 5C. The data are all consistent with $n_{CI}$ = 4. The traces are the average of three technical replicates, ± S.D. See Supplementary Figure 7 for supporting data on the *Pd*-SBP preparations and Materials and Methods for experimental conditions and protocols.

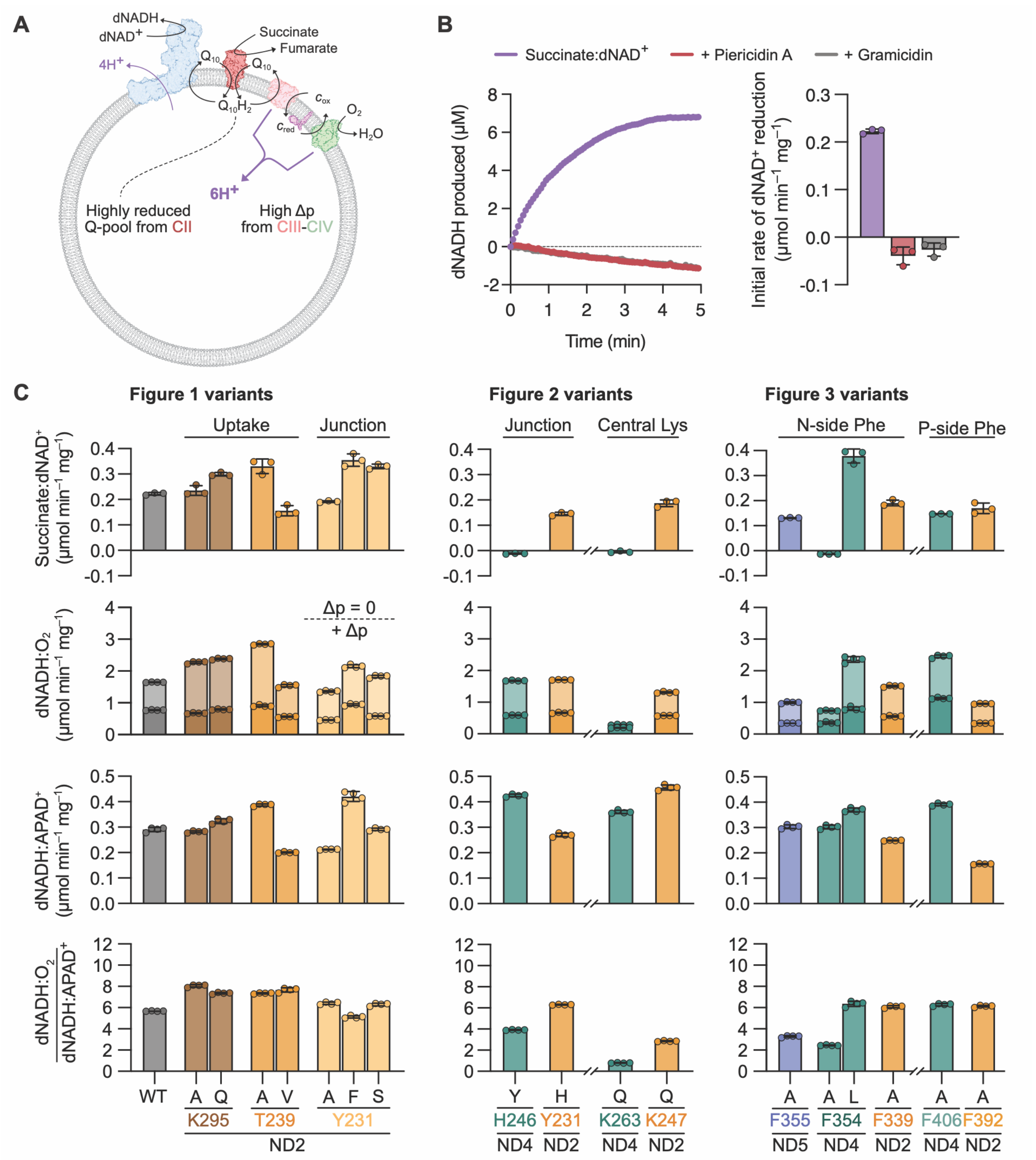


**Supplementary Figure 7. Reverse electron transfer (RET) in SBPs from *P. denitrificans*.** (**A**) Schematic diagram of RET (succinate:dNAD$^+$ oxidoreduction) in *P. denitrificans* SBPs. Succinate/CII reduces ubiquinone-10 ($Q_{10}$) to ubiquinol-10 ($Q_{10}H_2$), which is reoxidized by CIII-CIV (6 $H^+$ pumped per succinate) to establish Δp. Providing both Q-pool reduction and Δp are high, the combined free energy from 4 $H^+$ transferring back across the membrane through CI is sufficient to drive RET, reducing dNAD$^+$. (**B**) Experimental data on RET by WT *Pd*-SBPs. Rates of dNAD$^+$ reduction are measured at the start of the trace, which levels out as the system enters a steady state with dNAD$^+$ reduction by RET balanced by dNADH:$O_2$ oxidoreduction (CI-CIII-CIV). No RET is observed with piericidin A (CI inhibitor) or gramicidin (an 'uncoupler' which dissipates Δp). (**C**) Rates of RET and dNADH:$O_2$ oxidoreduction (± gramicidin) used to calculate the normalized rates of RET in Figures 1, 2 and 3 (n = 3, technical replicates, ± S.D.). The rate of RET depends on the *thermodynamic* driving force (the Q-pool status and Δp) as well as on the catalytic competance of the variant. The ability of the SPBs to maintain Δp is reflected in the ratio of dNADH:$O_2$ rates (± Δp). The rate is also expected to be *kinetically* related to the rate of 'forward' catalysis: rates of RET in Figures 1, 2 and 3 have been normalized by rates of dNADH:$O_2$ oxidoreduction (Δp = 0) and thus retain thermodynamic variations that preclude direct comparisons. For *Pd*-K263Q$^{ND4}$ the rates of RET and dNADH:$O_2$ oxidoreduction are very low so their ratio is not reported in Figure 2. Rates of dNADH:APAD$^+$ oxidoreduction are used as a proxy for the *Pd*-CI abundance, to normalize the rates of turnover (dNADH:$O_2$ + gramicidin divided by dNADH:APAD$^+$): the normalized data match the corresponding normalized data from membrane preparations presented in Figures 1, 2 and 3. See Materials and Methods for experimental conditions and protocols.

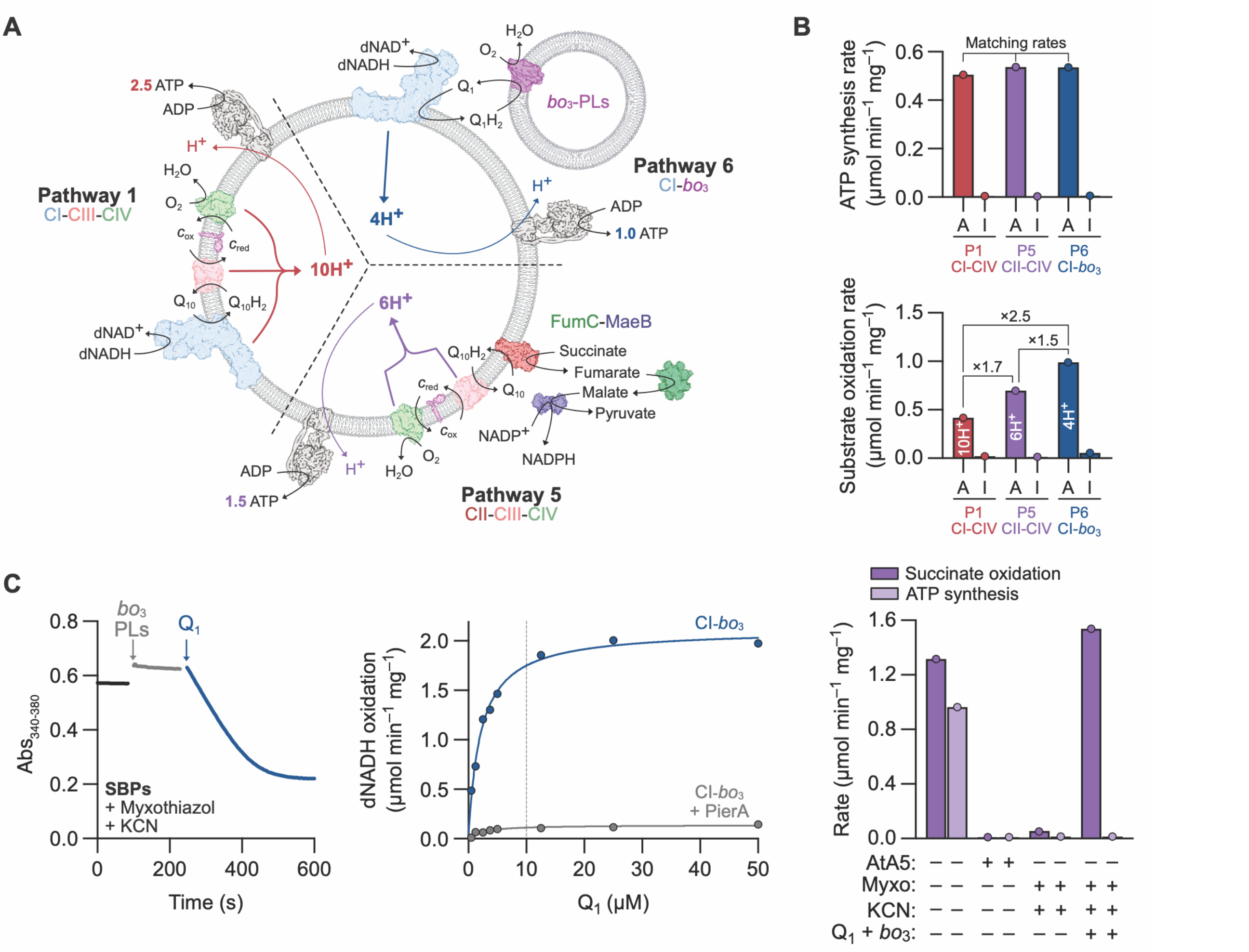


**Supplementary Figure 8. Principles of the *Pd*-CI stoichiometry measurements using SBPs from *P. denitrificans*.** (**A**) Proton-pumping by CI, CIII and CIV creates a Δp that is used by *P. denitrificans* ATP synthase to produce ATP (4 $H^+$ per ATP). Pathway 1 uses CI-CIII-CIV to pump 10 $H^+$ per dNADH oxidized. Pathway 5 uses CII-CIII-CIV to pump 6 $H^+$ per succinate oxidized, using a coupled detection assay (FumC-MaeB) to monitor CII activity as $NADP^+$ reduction (*58*). Pathway 6 (CIII and CIV inhibited) uses CI to pump 4 $H^+$ per dNADH oxidized; continuous catalysis is enabled by proteoliposomes containing cytochrome $bo_3$ oxidase ($bo_3$-PLs), together with low concentrations of a soluble quinone ($Q_1$) that equilibrates with the SBP $Q_{10}$ pool, forming $Q_1H_2$ that is reoxidized by the $bo_3$-PLs. This approach improves on the AOX addition used previously (*23*) as it avoids introducing detergents that impact membrane integrity. (**B**) Example data (for *Pd*-F406A$^{ND4}$) shows that, at matching rates of ATP synthesis, the rates of substrate oxidation are proportional to the number of protons pumped by each pathway. Substrate oxidation rates have been experimentally adjusted (marked A) using piericidin A (CI) or atpenin A5 (CII) inhibitors, or fully inhibited (marked I). The relative rates of substrate oxidation are used to calculate the stoichiometry of CI ($n_{CI}$, shown for $n_{CI}$ = 4), based on the ratios between them (2.5 = 10/4, 1.7 = 10/6, 1.5 = 6/4). Stoichiometry measurements in Figures 1, 2 and 3 display data over a range of substrate oxidation rates (rather than matching at a single rate of ATP synthesis) and determine $n_{CI}$ by plotting the normalized log of the sum of the squared errors (data vs. fit) at each candidate value of $n_{CI}$. The global best fit is at the minimum point of the plot. (**C**) Supporting data on Pathway 6. Left, 10 µM $Q_1$ enables sustained *Pd*-CI catalysis (dNADH oxidation monitored at 340 nm) by mediating between the *Pd*-SBPs $Q_{10}$-pool and the $bo_3$-PLs: in the presence of dNADH, SBPs treated with myxothiazol and KCN to inhibit CIII-CIV are inactive until both $bo_3$-PLs and $Q_1$ are added. The concentration of KCN used selectively inhibits CIV, not $bo_3$. Middle, dependence of the rate of Pathway 6 catalysis on $Q_1$ concentration in the presence and absence of piericidin A to inhibit *Pd*-CI. The vertical marker indicates the chosen concentration. Right, control data showing that $bo_3$-PLs themselves do not synthesize ATP. Pathway 5 catalysis (succinate oxidation and ATP synthesis) is sensitive to atpenin A5 (AtA5, CII inhibitor) and myxothiazol (Myxo) plus KCN (CIII-CIV inhibitors); addition of $bo_3$-PLs and $Q_1$ (to activate succinate oxidoreduction by CII-$bo_3$) restores succinate oxidation but not ATP synthesis. See Materials and Methods for experimental conditions and protocols.

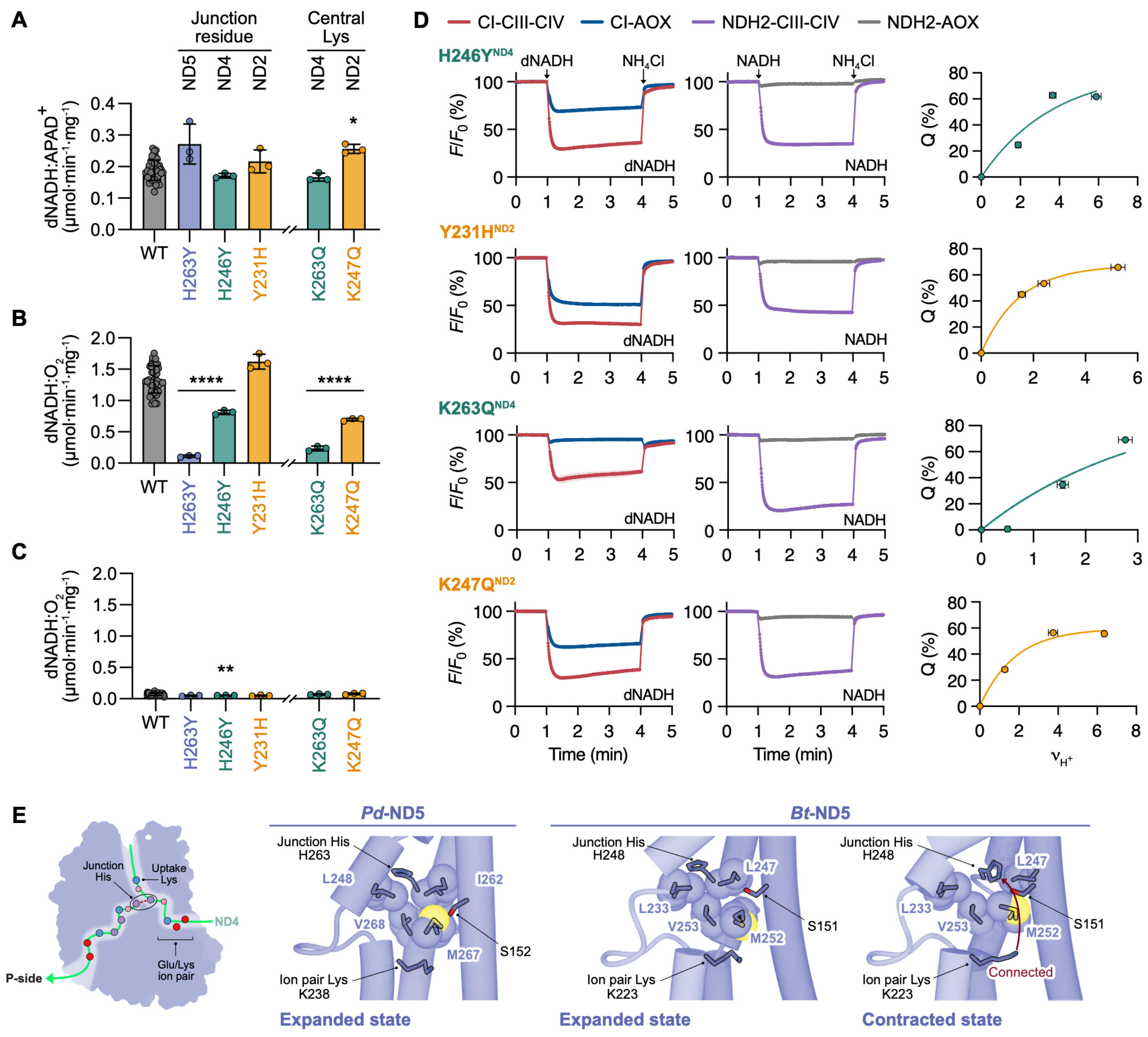


**Supplementary Figure 9. Supporting data and analyses for Figure 2.** Rates of dNADH:APAD$^+$ oxidoreduction (**A**) and dNADH:$O_2$ oxidoreduction without (**B**) and with (**C**) the CI inhibitor piericidin A (5 µM) were determined in independent *Pd*-membrane preparations ($N \geq 3$, ± S.D.). (**D**) Proton pumping by *Pd*-CI variants assessed by ACMA fluorescence quenching in *Pd*-SBPs. ACMA quenching was determined for the four proton-pumping pathways shown in Supplementary Figure 5A and analysed as illustrated in Supplementary Figure 5B-C. Supplementary Figure 5C shows corresponding data for WT *Pd*-CI. The traces shown are the average of three technical replicates, ± S.D. Data for *Pd*-H246Y$^{ND4}$ and *Pd*-K263Q$^{ND4}$ are inconsistent with a four-proton stoichiometry. However, the rate of dNADH oxidation by *Pd*-K263Q$^{ND4}$ is very slow (Supplementary Figure 7) precluding firm conclusions about proton pumping. See Supplementary Figure 7 for supporting data on the *Pd*-SBP preparations. See Materials and Methods for experimental conditions and protocols. (**E**) The peak route of the central axis in ND5. The cartoon shows the peak profile of the central axis and the uptake pathway viewed from the membrane (red – Glu, blue – Lys, purple – His, pink –Ser/Tyr/Thr); the flat pathway does not exist due to the absence of the central-Lys. The structures show the view along the central axis from the ion pair, showing TMH5, 7, 8 with van der Waals radii for relevant residues. The structure for *Pd*-ND5 (PDB: 8QBY) is in the expanded state, whereas two states have been observed for mammalian (*Bos taurus*) CI, expanded (PDB: 9R12) and contracted (PDB: 9R14) (*16*). In the contracted state the ion-pair Lys is connected to the junction-His by a hydrated pathway that includes *Pd*-S152$^{ND5}$ (the 'upward' part of the peak). In the expanded state the junction-His has moved away from *Pd*-S152$^{ND5}$ and is connected to the proton-output pathway (the 'downward' part of the peak). The disconnection is assisted by movement of *Pd*-M267$^{ND5}$ (yellow) into the hydrated region.

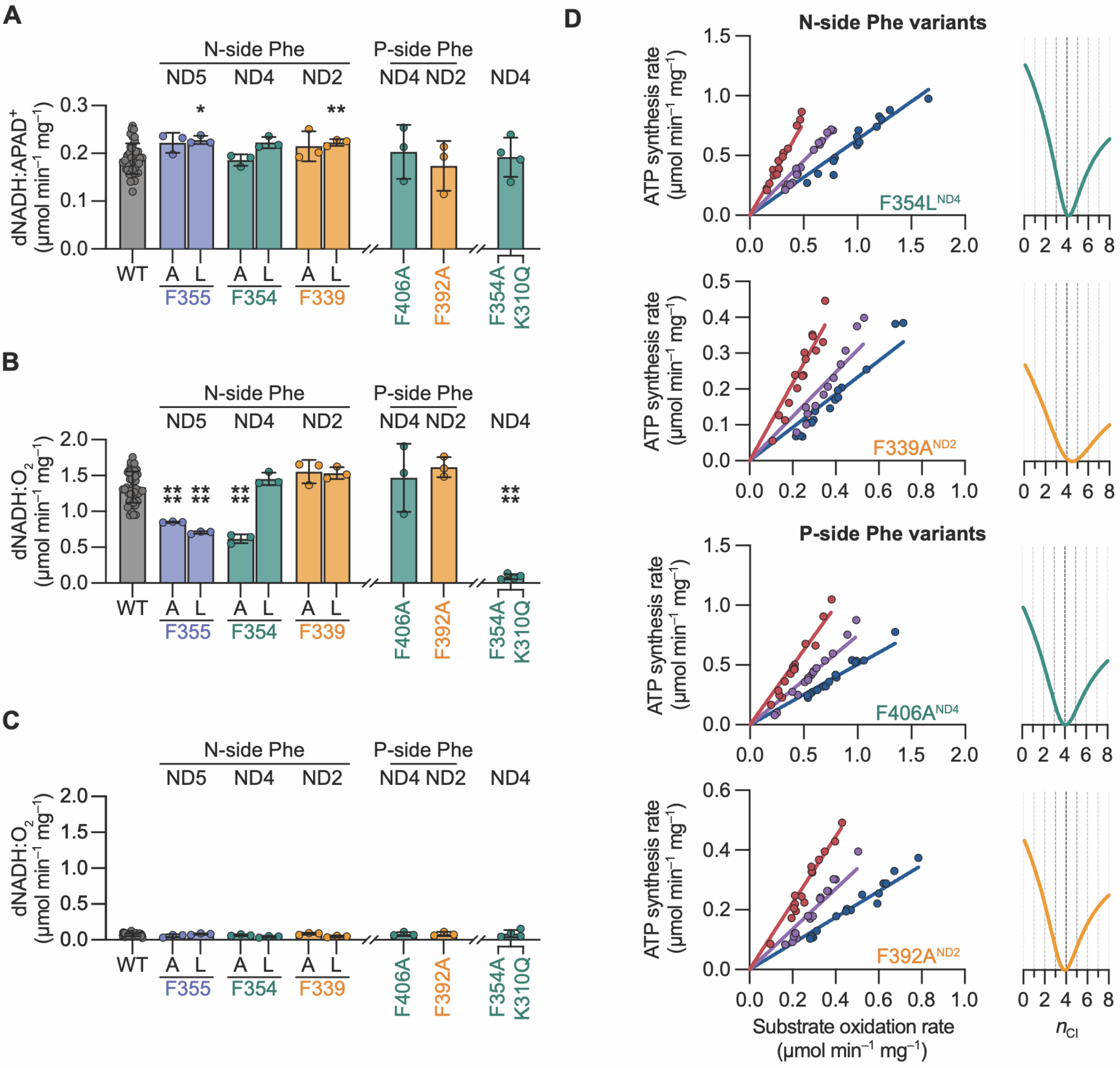


**Supplementary Figure 10. Supporting data and analyses for Figure 3.** Rates of dNADH:APAD$^+$ oxidoreduction (**A**) and dNADH:$O_2$ oxidoreduction without (**B**) and with (**C**) the CI inhibitor piericidin A (5 μM) were determined in independent *Pd*-membrane preparations ($N \geq 3$, ± S.D.). F354:K310$^{ND4}$ is a double variant. (**D**) Determination of the proton-pumping stoichiometries of the N-side Phe variants *Pd*-F354L$^{ND4}$ and *Pd*-F339A$^{ND2}$ and the P-side Phe variants *Pd*-F406$^{ND4}$ and *Pd*-F392A$^{ND2}$ (see Supplementary Figure 8 for further information). Left: ATP synthesis rates plotted against substrate oxidation rates for three pathways (dNADH:$O_2$ by CI-CIII-CIV, succinate:$O_2$ by CII-CIII-CIV, and dNADH:$O_2$ by CI-*bo*$_3$, using *bo*$_3$-proteoliposomes). The lines shown are the global best fits, and their relative gradients reflect the pathway stoichiometries. Right: The *Pd*-CI stoichiometry ($n_{CI}$) is determined by plotting the normalized log of the sum of the squared errors (data vs. fit) at each candidate value of $n_{CI}$. The global best fit is at the minimum point (*Pd*-F354L$^{ND4}$ = 4.17, $R^2$ = 0.932; *Pd*-F339A$^{ND2}$ = 4.52, $R^2$ = 0.873; *Pd*-F406A$^{ND4}$ = 4.06, $R^2$ = 0.923; *Pd*-F392A$^{ND2}$ = 3.88, $R^2$ = 0.934) and the y-scale is from 1 to 100. See Materials and Methods for experimental conditions and protocols.

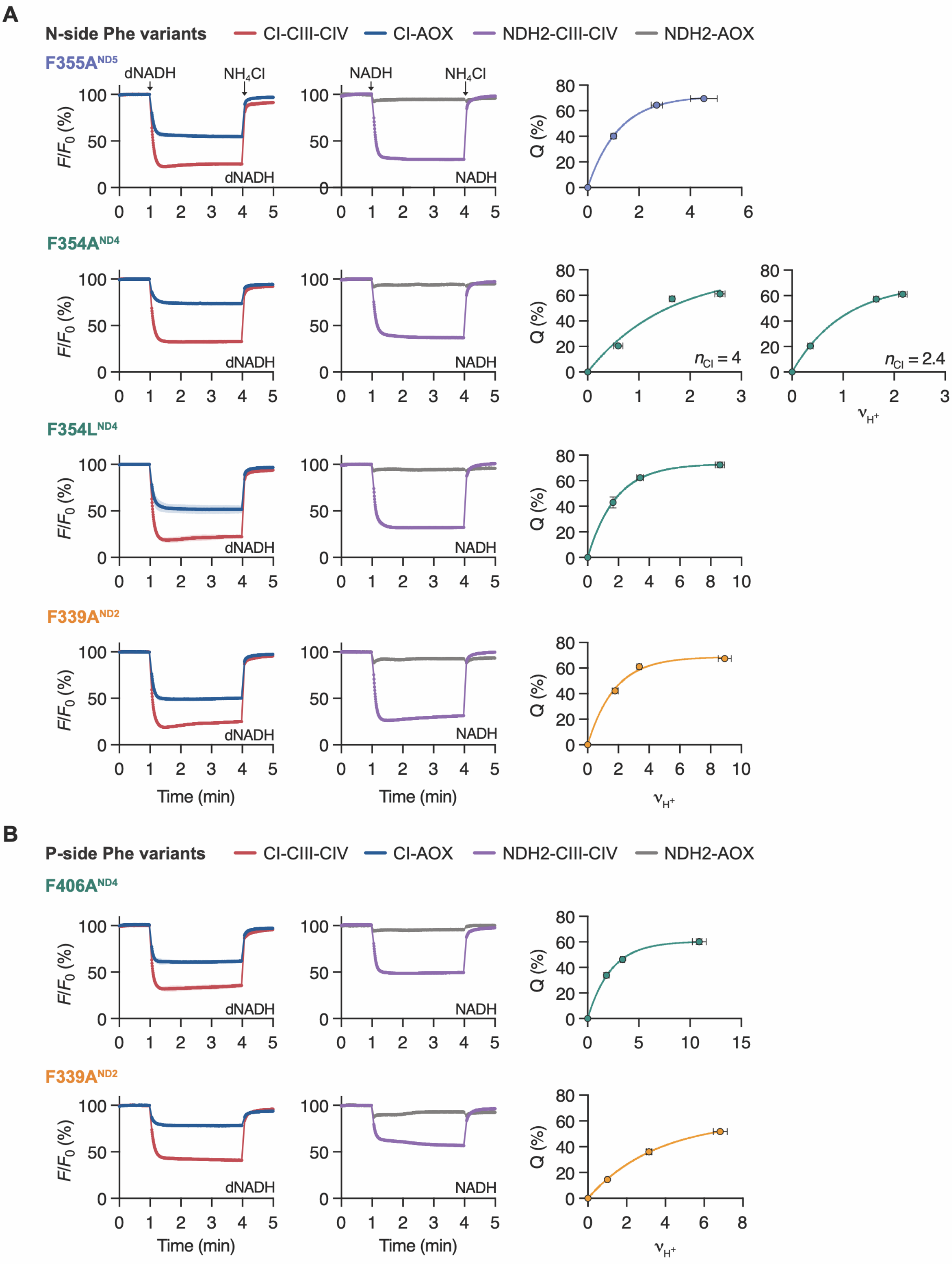


**Supplementary Figure 11. Proton pumping by the N- and P-side Phe variants of *Pd*-CI in Figure 3 assessed by ACMA fluorescence quenching in *Pd*-SBPs.** ACMA quenching was determined for the four proton-pumping pathways shown in Supplementary Figure 5A and analysed as illustrated in Supplementary Figure 5B-C for N-side (**A**) and P-side (**B**) Phe variants. Supplementary Figure 5C shows corresponding data for WT *Pd*-CI. The data fit for *Pd*-F354A$^{ND4}$ using a four-proton stoichiometry is poor so an alternative fit using the $n_{CI}$ value of 2.4 from the direct stoichiometry measurement (Figure 3D) is also shown. The traces are the average of three technical replicates, ± S.D. See Supplementary Figure 7 for supporting data on the *Pd*-SBP preparations and Materials and Methods for experimental conditions and protocols.

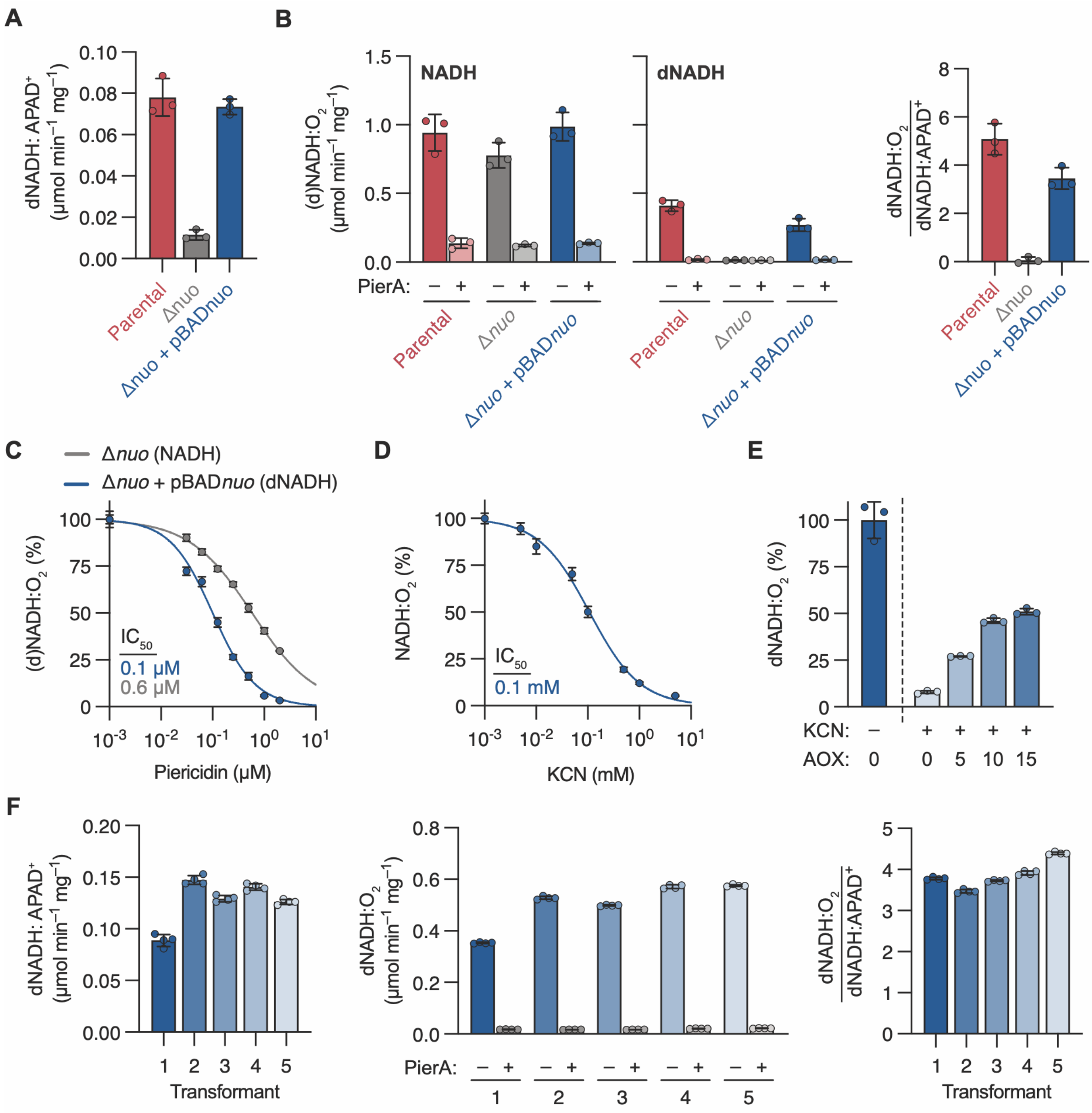


**Supplementary Figure 12. Expression of wild-type *E. coli* complex I in the Δ*nuo E. coli* strain.** (**A**) Rates of dNADH:APAD$^+$ oxidoreduction in *E. coli* membranes from the parental strain, the Δ*nuo* (*Ec*-CI operon knockout) strain, and the Δ*nuo* strain complemented with the pBAD*nuo* plasmid expressing *Ec*-CI. (**B**) Rates of (d)NADH:$O_2$ oxidoreduction in *E. coli* membranes from the three strains. All three show activity with NADH (left) due to native expression of the alternative NADH:quinone oxidoreductase NDH2, whereas dNADH activities (middle) are specific to CI. Rates are also shown in the presence of 2 μM piericidin A, a canonical CI inhibitor. The piericidin A inhibits dNADH oxidation by *Ec*-CI completely, but was found to partially inhibit NDH2 catalysis also (reflected in the inhbition of NADH oxidation, see also panel C). Right, dNADH:$O_2$ divided by dNADH:APAD$^+$ to normalize the rate for *Ec*-CI expression. Data in panels (**A**) and (**B**) are from three independent preparations (N = 3, ± S.D). (**C**) Separate titrations for the inhibition of *Ec*-CI (Δ*nuo* + pBAD*nuo*, dNADH:$O_2$) and NDH2 (Δ*nuo*, NADH:$O_2$) by piericidin A in *E. coli* membranes, showing that piericidin A-inhibition is not selective. (**D**) Titration of the inhibition of NADH:$O_2$ oxidoreduction by KCN in *E. coli* membranes (Δ*nuo* + pBAD*nuo*), which inhibits the *E. coli* quinol oxidases (expression of the cytochrome $bo_3$ oxidase, $bo_3$, is expected to dominate under the cell culture conditions used) (*65*). (**E**) Addition of exogenous AOX restores dNADH:$O_2$ activity in KCN-inhibited *E. coli* membranes (10 μg mL$^{-1}$ membranes, 2.5 mM KCN, AOX as indicated in μg mL$^{-1}$). (**F**) Rates of dNADH:APAD$^+$ oxidoreduction (left) show variable expression of WT *Ec*-CI in five independent pBAD*nuo* transformants (numbered 1 to 5). Corresponding rates of dNADH:$O_2$ oxidoreduction (in the presence and absence of 2 μM piericidin A, middle) follow the same pattern, but differences in expression between the transformants do not affect the normalized rate (right). Data in panels (**C-F**) are technical replicates (n = 3, ± S.D). See Materials and Methods for experimental conditions and protocols.

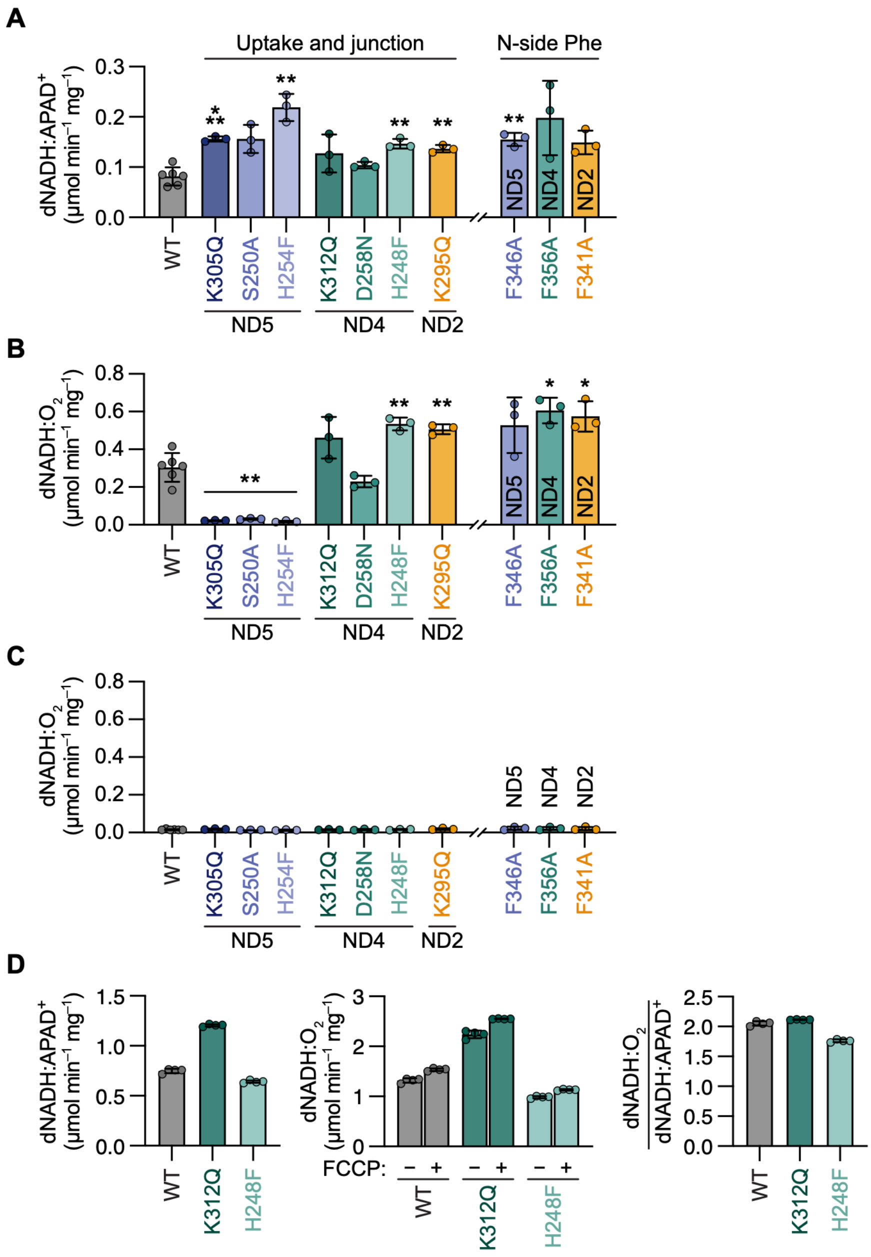


**Supplementary Figure 13. Supporting catalytic activity data on the *E. coli* complex I variants in Figure 4.** Rates of dNADH:APAD$^+$ oxidoreduction (**A**) and dNADH:$O_2$ oxidoreduction without (**B**) and with (**C**) the CI inhibitor piericidin A (5 µM) were determined in independent *Ec*-membrane preparations ($N \geq 3$, ± S.D.). (**D**) Rates of dNADH:APAD$^+$ (left) and dNADH:$O_2$ (middle) catalysis in SBPs from *E. coli* ($n$ = 3 technical replicates, ± S.D.). dNADH:$O_2$ measurements were performed in the presence and absence of 2 µM FCCP, a protonophore added to dissipate Δp. Rates in the presence of FCCP (Δp = 0) were normalized by the dNADH:APAD$^+$ measurements (right). Note that the data in panel D were measured at pH 7.5 for comparison with data from ACMA measurements. See Materials and Methods for experimental conditions and protocols.

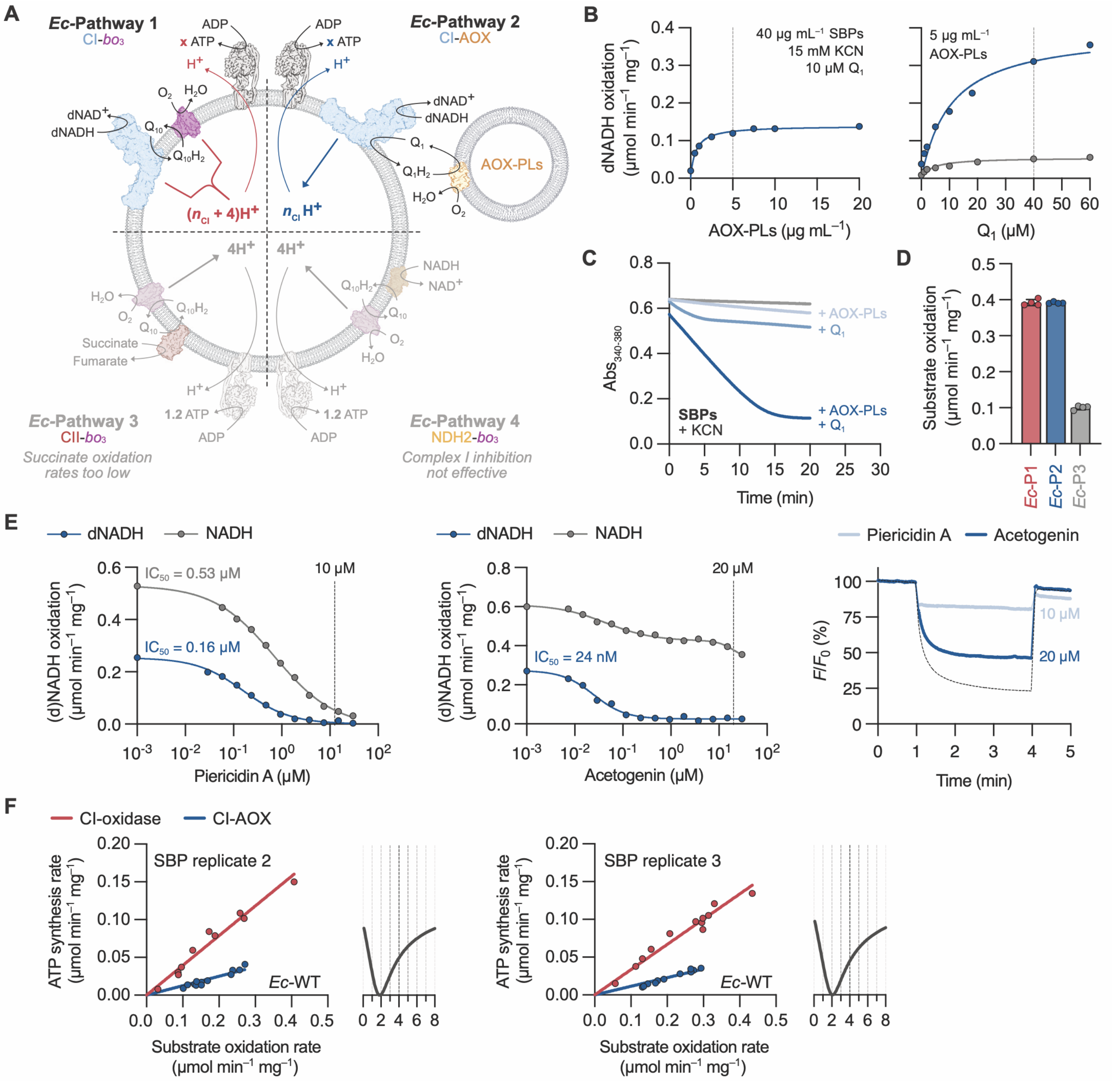


**Supplementary Figure 14. Optimization of stoichiometry measurements using SBPs from *E. coli*.** (**A**) Two-pathway measurement for the proton-pumping stoichiometry of *Ec*-CI based on the system used for *Pd*-CI (Supplementary Figure 7). Proton-pumping by CI and cytochrome *bo*$_3$ oxidase creates a Δp that is used by *E. coli* ATP synthase to produce ATP (3.3 H$^+$ per ATP). *Ec*-Pathway 1 uses CI-*bo*$_3$ to pump ($n_{CI}$ + 4) H$^+$ per dNADH oxidized (*39*, *40*). *Ec*-Pathway 2 (*bo*$_3$ inhibited) uses CI to pump $n_{CI}$ H$^+$ per dNADH oxidized. Due to the presence of *bo*$_3$ in the *Ec*-SBPs, continuous catalysis is enabled by proteoliposomes containing the alternative oxidase (AOX-PLs), mediated by ubiquinone-1 ($Q_1$). AOX-PLs are cyanide insensitive allowing specific inhibition of the native *bo*$_3$. *Ec*-Pathways 1 and 2 were used to estimate $n_{CI}$. *Ec*-Pathway 3 (CII-*bo*$_3$, 4 H$^+$ per succinate oxidized) and *Ec*-Pathway 4 (NDH2-*bo*$_3$, 4 H$^+$ per NADH oxidized) were unsuccessful as candidate pathways. (**B**) Supporting data on *Ec*-Pathway 2. Left, the maximum rate of dNADH oxidation is achieved at 5 μg mL$^{-1}$ AOX-PLs. Right, 40 μM $Q_1$ provides a sufficiently high rate for stoichiometry measurements. (**C**) Sustained catalysis by KCN-inhibited *Ec*-SBPs requires both $Q_1$ and AOX-PLs (concentrations as determined in **B**). (**D**) *Ec*-Pathway 3 (CII-*bo*$_3$) is too slow for stoichiometry measurements (n = 4 technical replicates, ± S.D). (**E**) For *Ec*-Pathway 4, a CI inhibitor capable of isolating NDH2 catalysis in *E. coli* has not been identified. Left, piericidin shows strong inhibition of CI (dNADH) but the high concentrations required also inhibit NDH2 (NADH). Middle, CI inhibition by an acetogenin (*72*) in *Ec*-SBPs shows no significant effect on NDH2 activity (NADH) but does not achieve complete CI inhibition. Right, ACMA traces recorded at the marked concentrations of maximal CI inhibition for piericidin A and the acetogenin. A substantial quench remains in the presence of the acetogenin. An uninhibited trace (dashed) is shown for comparison. (**F**) Stoichiometry measurements on independent *Ec*-SBP preparations (parental strain), matching the measurement and analyses shown in Figure 4D. Best-fit values for $n_{CI}$ were: replicate 2 = 1.86 ($R^2$ = 0.975) and replicate 3 = 2.08 ($R^2$ = 0.979).

## ND5

| | | Uptake | | Junct. | N-Phe | Ion pair | Central | Terminal | P-Asp |
|---|---|---|---|---|---|---|---|---|---|
| Bacteria | *P. denitrificans* | K314 | S259 | H263 | F355 | K238 | - | K413 | D414 |
| | *E. coli* | K305 | S250 | H254 | F346 | K229 | - | K399 | D400 |
| | *T. thermophilus* | K292 | S237 | H241 | F333 | K216 | - | K385 | D386 |
| | *M. smegmatis* | K307 | S252 | H256 | F350 | K231 | - | K402 | D403 |
| Plants | *V. radiata* | K308 | S253 | H257 | F349 | K232 | - | K401 | D402 |
| | *A. thaliana* | K308 | S253 | H257 | F349 | K232 | - | K401 | D402 |
| Yeasts | *P. pastoris* | K299 | S244 | H248 | F340 | K223 | - | K393 | D394 |
| | *C. thermophilum* | K299 | S244 | H248 | F340 | K223 | - | K392 | D393 |
| | *Y. lipolytica* | K302 | S247 | H251 | F343 | K226 | - | K396 | D397 |
| Ciliates | *Polytomella* Sp. | K288 | S239 | H243 | F331 | K218 | - | K374 | E375 |
| | *T. thermophila* | K481 | S426 | H430 | F522 | K405 | - | K574 | H575 |
| Insects | *D. melanogaster* | K266 | S212 | H216 | F307 | K191 | - | K359 | D360 |
| Mammals | *S. scrofa* | K299 | S244 | H248 | F340 | K223 | - | K392 | D393 |
| | *O. aries* | K299 | S244 | H248 | F340 | K223 | - | K392 | D393 |
| | *M. musculus* | K299 | S244 | H248 | F340 | K223 | - | K392 | D393 |
| | *B. taurus* | K299 | S244 | H248 | F340 | K223 | - | K392 | D393 |
| | *H. sapiens* | K299 | S244 | H248 | F340 | K223 | - | K392 | D393 |

## ND4

| | | Uptake | | Junct. | N-Phe | Ion pair | Central | Terminal | P-Phe |
|---|---|---|---|---|---|---|---|---|---|
| Bacteria | *P. denitrificans* | K310 | S254 | H246 | F354 | K232 | K263 | E405 | F406 |
| | *E. coli* | K312 | D258[1] | H248 | F356 | K234 | K265 | E407 | F408 |
| | *T. thermophilus* | K282 | D228[1] | H218 | F326 | K204 | K235 | E377 | F378 |
| | *M. smegmatis* | M321 | A265 | A257 | F365 | K243 | K274 | E416 | F417 |
| Plants | *V. radiata* | K305 | S249 | H241 | F349 | K227 | K258 | E400 | F401 |
| | *A. thaliana* | K305 | S249 | H241 | F349 | K227 | K258 | E400 | F401 |
| Yeasts | *P. pastoris* | K302 | S246 | H238 | F346 | K224 | K255 | E398 | F399 |
| | *C. thermophilum* | K340 | S284 | H276 | F384 | K262 | K293 | E437 | F438 |
| | *Y. lipolytica* | K299 | S243 | H235 | F343 | K221 | K252 | E395 | F396 |
| Ciliates | *Polytomella* Sp. | K273 | S217 | H209 | F317 | K195 | K226 | E368 | V369 |
| | *T. thermophila* | K331 | S278 | H270 | F375 | K256 | K287 | E426 | F427 |
| Insects | *D. melanogaster* | K271 | S216 | H208 | F315 | K194 | K225 | E366 | I367 |
| Mammals | *S. scrofa* | K283 | S228 | H220 | F327 | K206 | K237 | E378 | L379 |
| | *O. aries* | K283 | S228 | H220 | F327 | K206 | K237 | E378 | L379 |
| | *M. musculus* | K283 | S228 | H220 | F327 | K206 | K237 | E378 | L379 |
| | *B. taurus* | K283 | S228 | H220 | F327 | K206 | K237 | E378 | L379 |
| | *H. sapiens* | K283 | S228 | H220 | F327 | K206 | K237 | E378 | L379 |

| ND2 | | | | | | | | | |
|---|---|---|---|---|---|---|---|---|---|
| | | **Uptake** | | **Junct.** | **N-Phe** | **Ion pair** | **Central** | **Terminal** | **P-Phe** |
| Bacteria | *P. denitrificans* | K295 | T239 | Y231 | F339 | K217 | K247 | K391 | F392 |
| | *E. coli* | K295 | S239 | Y231 | F341 | K217 | K247 | K395 | F396 |
| | *T. thermophilus* | K256 | V208 | Y200 | F296 | K186 | K216 | K345 | Y346 |
| | *M. smegmatis* | K335 | T279 | Y271 | F379 | K257 | K287 | K432 | F433 |
| Plants | *V. radiata* | K306 | T251 | Y243 | F349 | K229 | K259 | K399 | F400 |
| | *A. thaliana* | K316 | T262 | Y254 | F360 | K240 | K270 | K410 | F411 |
| Yeasts | *P. pastoris* | K289 | T233 | Y225 | F333 | K211 | K241 | K416 | Y417 |
| | *C. thermophilum* | K337 | T276 | Y268 | F381 | K254 | K284 | K451 | Q452 |
| | *Y. lipolytica* | K282 | T233 | Y225 | F324 | K211 | K241 | K383 | L384 |
| Ciliates | *Polytomella* Sp. | R235 | A189 | F181 | W276 | K167 | K197 | K303 | A304 |
| | *T. thermophila*[2] | K337 | I281 | Y273 | V31-B | K259 | F289[3] | K85-B | F86-B |
| Insects | *D. melanogaster* | R172 | A126 | M118 | T212 | K104 | K134 | K259 | W260 |
| Mammals | *S. scrofa* | R176 | G127 | T119 | F216 | K105 | K135 | K263 | W264 |
| | *O. aries* | R176 | G127 | T119 | F216 | K105 | K135 | K263 | W264 |
| | *M. musculus* | R176 | G127 | T119 | F216 | K105 | K135 | K263 | W264 |
| | *B. taurus* | R176 | G127 | T119 | F216 | K105 | K135 | K263 | W264 |
| | *H. sapiens* | R176 | G127 | T119 | F216 | K105 | K135 | K263 | W264 |

**Supplementary Table 1. Sequence numbers and key residues in all species of complex I for which high-resolution structures (<3.5 Å) have been determined.** Sequence alignments are shown in Supplementary Figure 2. The structure coordinate files are: *Paracoccus denitrificans*, PDB:8QBY (*14*); *Escherichia coli*, PDB:7Z7S (*6*); *Thermus thermophilus*, PDB:4HEA (*37*); *Mycobacterium smegmatis*, PDB:8E9G (*33*); *Vigna radiata*, PDB:8E73 (*73*); *Arabidopsis thaliana*, PDB:8BPX (*12*); *Pichia pastoris*, PDB:9IHR (*74*); *Chaetomium thermophilum*, PDB:7ZMG (*75*); *Yarrowia lipolytica*, PDB:6YJ4 (*76*); *Polytomella* Sp., PDB:7ARD (*77*); *Tetrahymena thermophila*, PDB:7TGH (*69*); *Drosophila melanogaster*, PDB:8B9Z (*78*); *Sus scrofa*, PDB:8UD1 (*15*); *Ovis aries*, PDB:6ZKD (*5*); *Mus musculus*, PDB:8OM1 (*13*); *Bos taurus*, PDB:8Q48 (*31*); *Homo sapiens*, PDB:9TI4 (*79*).

Footnotes:

[1]*Ec*-Asp258$^{ND4}$ and *T. thermophilus* Asp228$^{ND4}$ are spatially conserved with the uptake-Ser but not conserved in sequence (see Supplementary Figure 2).

[2]*T. thermophila* ND2 is split into two proteins (Ymf65/ND2A and ND2B) as shown in Supplementary Figure 2.

[3]In *T. thermophila*, Phe289$^{ND2A}$ is found in the position of the central-Lys, but the ε-amine of Lys269$^{ND2A}$ is nearby and may be able to functionally substitute for it.
